# Competing Molecular Interactions Govern the Dynamical Arrest of G3BP1 Condensates

**DOI:** 10.64898/2026.07.31.741769

**Authors:** Anurag Singh, Vicky Liu, Tharun Selvam Mahendran, Aoon Rizvi, Bhargavi Gindra, Brian Freibaum, Hong Joo Kim, J. Paul Taylor, Rohit V. Pappu, Priya R. Banerjee

## Abstract

G3BP1 is a central scaffold of stress granules (SGs). Upon cellular stress, G3BP1 forms complex coacervates with translationally repressed mRNAs and recruits multiple RNA-binding proteins to form reversible biomolecular condensates. Persistent SGs are linked to age-dependent dynamical arrest and impaired disassembly. Here, we employ active and passive nanoscale rheology with optical tweezers to show that G3BP1 condensates evolve from being dominantly viscous fluids to dynamically arrested network glasses characterized by nanoscale caging and elastic memory. Integrating atomistic and coarse-grained simulations with experiments, we find that electrostatic interactions between the oppositely charged intrinsically disordered regions drive condensate ageing. RNA modulates these interactions in a length-and structure-dependent manner, delaying dynamic arrest, whereas Caprin-1 binding to the NTF2L domain has little effect. Together, these findings reveal how competing inter-IDR and IDR–RNA interactions govern condensate ageing and material-state transitions. The findings have broader implications for the regulation of SG dynamics in cells.

## Introduction

Biomolecular condensates formed through associative phase separation such as complex coacervation of proteins and nucleic acids have been proposed to drive sub-cellular organization^1–8^. Cytoplasmic stress granules (SGs) are multicomponent condensates that assemble in response to a multitude of cellular stresses^9–14^. Under stress, mRNAs released from stalled translational machinery interact with cytoplasmic ribonucleoproteins to undergo phase separation, leading to the formation of SGs^15^. When the stress is released, SGs undergo disassembly through ATP-dependent active processes orchestrated by RNA and protein chaperones^16,17^. However, under prolonged stress and/or in the presence of RNA-binding proteins (RBPs) carrying disease-associated mutations, SG disassembly can be impaired, leading to persistent SGs^18,19^. These persistent granules are thought to possess different material properties when compared to nascent SGs and they have the potential to evolve into cytotoxic inclusions linked to neurodegenerative diseases^20,21^.

Understanding the physical principles and the biological factors that govern the temporal evolution of material properties of SGs is central to establishing a mechanistic role of SGs in neurodegenerative diseases^22^. Several orthogonal models have been proposed to account for the ageing of biomolecular condensates. Pathological SG ageing is often attributed to amyloid formation by aggregation-prone RBPs^21,23–25^, such as TDP-43, hnRNPA1, and FUS that localize to SGs^26^ (**Supplementary Fig. 1**). Emerging evidence suggests that the condensate interior can suppress amyloid assembly^25^. In this scenario, amyloid growth occurs predominantly in the coexisting dilute phase, whereas condensate interfaces promote nucleation^24,25,27^. Another emerging mechanism is the oxidative stress-induced de-mixing and aggregation of RBPs, such as TDP-43, within stress granules^23^. Growing evidence also suggests that condensates can undergo dynamical arrest through material-state transitions. These transitions appear to be independent of amyloid formation and they include the formation of ordered viscoelastic solids^28^ and glass-like states^29,30^. Here, we explore the molecular driving forces that underlie non-amyloid ageing pathways using condensates formed by G3BP1 as a model system.

G3BP1, a multi-domain RBP, is the core component of the protein-RNA interaction network of SGs^31–33^. Upon cellular stress, G3BP1 forms complex coacervates with translationally repressed mRNAs and recruits multiple RNA-binding proteins to assemble reversible biomolecular condensates^15,32,33^. We investigated the ageing dynamics of condensates formed by full-length G3BP1 in the absence of RNA and compare them with those of G3BP1–RNA complex coacervates formed in the presence of RNA. Complex coacervation^34^ is also known to be modulated by protein binding partners that function as ligands that either enhance or suppress phase separation^15,32,33^. We therefore examined the effect of Caprin-1, a well-established G3BP1 binding partner with a central role in stress granule assembly^35,36^, on the ageing dynamics of G3BP1 condensates.

Employing quantitative micro and nano-scale rheology, in parallel with simulations and complementary biophysical measurements, we discovered that condensates that form via crowder-induced depletion-mediated attractions^37^ undergo ageing-dependent dynamical arrest. This arrest is the result of a transition to a viscoelastic network glass, characterized by the formation of nanoscale cages. RNA binding via the RNA recognition motif (RRM) and the R/G-rich intrinsically disordered region 3 (IDR3) fluidizes G3BP1 condensates and enhances the barrier for dynamical arrest through molecular frustration. In contrast, heterotypic protein-protein interactions that occur through the NTF2L domain of G3BP1^38,39^ do not impact their intrinsic dynamics. Our findings suggest that the competing network of RNA-protein interactions in SGs provides a mechanism for tuning condensate fluidity, with distinct RNA and protein partners playing differential roles in G3BP1-driven interactions that underlie condensate dynamical arrest as a function of ageing.

## Results and Discussion

### G3BP1 condensates undergo physical ageing without amyloid formation

G3BP1 is composed of an NTF2L dimerization domain that is also a protein recognition domain, an RRM, and three IDRs: IDR1 is mostly acidic, IDR2 contains proline-rich motifs, and IDR3 is composed of an R/G-rich region that binds RNA **(Fig. 1a)**. These sequence features allow G3BP1 to interact with RNA through the RRM and IDR3^40^ and protein partners such as Caprin-1 through the NTF2L domain^38^. In the absence of an interacting partner, G3BP1 populates a compact, closed conformation^31,32^, owing to the interactions mediated by the autoinhibitory electrostatic interactions between IDR1 and IDR3. These interactions also drive G3BP1 to undergo phase separation in vitro in the presence of molecular crowders^32^ in physiologically relevant buffer conditions.

**Figure 1.**
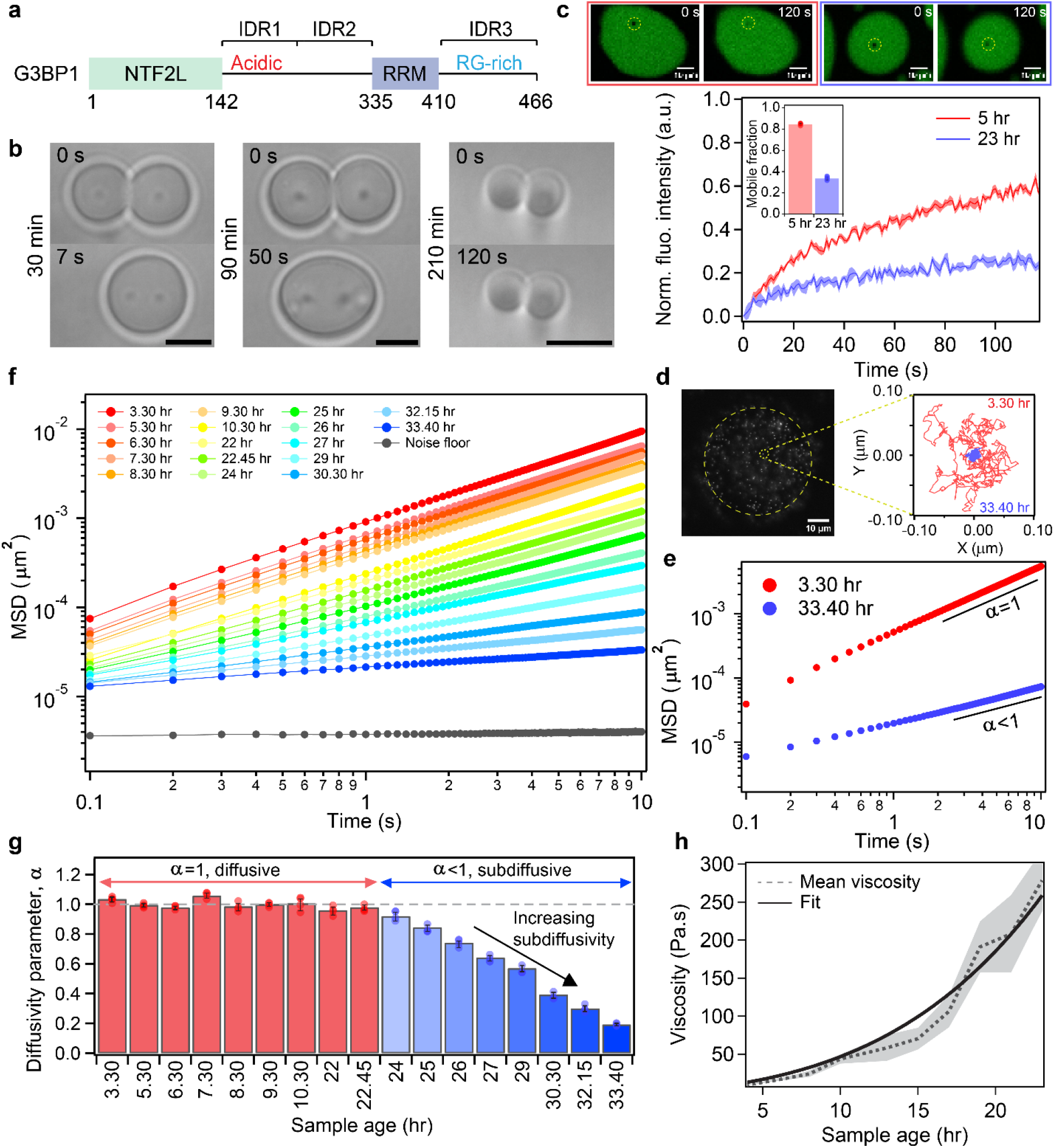
Homotypic G3BP1 condensates are metastable and undergo physical ageing. **a**, Schematic of G3BP1 with different interaction domains. **b**, Optical-tweezer-induced fusion of G3BP1 droplets at different time points of ageing. **c**, FRAP of Atto488-labeled G3BP1 in the dense phase: Top panel shows condensate (5 hr age, left; 23 hr age, right) images during a FRAP experiment at 0 s and 120 s time points. Bottom panel shows the fluorescence recovery curves. The inset shows the mobile fractions for 5 hr and 23 hr old condensates. **d**, Left panel: An epi-fluorescence image of G3BP1 condensates with fluorescently labeled 200-nm probe particles. Right panel: Representative trajectories of a probe particle in a G3BP1 condensate at 3:30 hr and 33:40 hr of age. **e**, Ensemble-average mean square displacement (MSD) of the probe particles in G3BP1 condensates at 3:30 hr and 33:40 hr of sample age. The slope of the MSDs at longer timescales characterizes the diffusion of the beads, with α = 1 corresponding to normal diffusion, and α < 1 corresponding to sub-diffusive behavior. **f**, Ensemble-average MSD of the probe particles at various condensate ages. The MSDs for which α = 1 holds at longer timescales were used to estimate the viscosity of G3BP1 condensates using the Stokes-Einstein relation. **g**, Variation of the diffusivity parameter, α, as a function of sample age. **h**, Viscosities are estimated from the MSDs for which α = 1 is shown as a function of condensate age. The dotted black line represents the mean value of viscosity, and the solid black line represents the fit using the equation 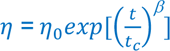. All measurements were repeated using at least three independently prepared samples.

We used crowder-induced G3BP1 condensates (50 µM G3BP1 in a buffer containing 10% Ficoll-70, 50 mM HEPES, 150 mM NaCl, pH 7.5)^32^ as the starting point of our study (concentration, crowder, and salt-dependent phase separation of G3BP1 is shown in **Supplementary Fig. 2**). To confirm that Ficoll-70 drives G3BP1 phase separation via depletion-mediated interactions^41^, we measured the partitioning of Fluorescein-labeled Ficoll-70 inside G3BP1 condensates at various concentrations of Ficoll-70 in the range 7.5% to 25% **(Supplementary Fig. 3)**. At all the tested concentrations, we observed exclusion of Ficoll-70 from G3BP1 condensates (partition coefficient < 1.0). Next, to characterize the material properties of G3BP1 condensates, we employed an optical-tweezer-based droplet fusion assay^42^ **(Fig. 1b)**. We observed three different stages of droplet fusion dynamics depending on the age of these condensates. At 30 minutes, the G3BP1 condensates were able to undergo rapid fusion and relaxation into a spherical shape, showing material properties dominated by the interfacial tension. However, after 90 minutes, G3BP1 condensates were unable to undergo complete shape relaxation into a spherical shape after fusion. The fused condensates instead took on shapes that were ellipsoidal, indicating a competition between viscosity and elasticity^43^ that frustrates droplet relaxation dynamics. After 210 minutes, G3BP1 droplets were unable to undergo fusion, suggesting that they were dynamically arrested. The observations of physical ageing of G3BP1 condensates were further confirmed using fluorescence recovery after photobleaching (FRAP) **(Fig. 1c, Supplementary Fig. 4)**. We observed ∼80% fluorescence recovery of the Atto488-labeled G3BP1 in the condensed phase at 5 hr of sample age, which markedly diminished to ∼30% after 23 hr of sample age **(Fig. 1c, inset)**. To test whether G3BP1 condensate ageing accompany amyloid-like fibril formation, as reported for canonical amyloid-forming proteins^24,25,27^, we employed Thioflavin-T (ThT) fluorescence (**Supplementary Fig. 5**). Utilizing microtubule-associated protein (MAPT) Tau condensates as a positive control^44^, we established that both nascent and aged G3BP1 condensates do not show any measurable ThT signal, indicating the absence of *bona fide* amyloid fibrils in G3BP1 condensates (**Supplementary Fig. 5b**). Together, these experiments indicate that G3BP1 condensates undergo physical ageing through a non-amyloid pathway.

Next, we used video particle tracking (VPT)-based nanorheology assay to probe G3BP1 condensate ageing with high spatial and temporal resolution^28,45–47^. For VPT assays, 200 nm-sized yellow-green fluorescence beads were embedded inside the G3BP1 condensates (**Fig. 1d**, left panel). Bead motion within the condensates was tracked to extract individual bead trajectories. Representative trajectories of a single bead at 3.5 hours and 33.5 hours of sample age are shown in **Fig. 1d** (right panel). The spatial extent of bead motion is markedly reduced at 33.5 hours, indicating dynamical arrest on the nanoscale within aged condensates. Trajectories from all the beads were utilized to estimate the ensemble-average mean-squared displacement (MSD). A progressive shift of the MSD to a lower value with condensate age indicates a reduced mobility of the beads (**Fig. 1e, 1f**). As the condensates aged, the MSDs continuously decreased, revealing progressively restricted bead mobility. We used the slope of the MSDs at longer timescales to characterize the motions of beads inside condensates, with α = 1 corresponding to normal diffusion and α < 1 corresponding to sub-diffusive behavior. Analysis of the ensemble average MSDs revealed the onset of sub-diffusive behavior around 24 hr of age (**Fig. 1g**), progressing to near dynamical arrest by ∼34 hr. The onset of sub-diffusivity is suggestive of the formation of nanoscale cages with sample age^28,48^.

From the MSD profiles for which α = 1 (up to ∼22.5 hours of sample age), we estimated the zero shear, bulk viscosity of G3BP1 condensates (**Fig. 1h**). We observe that condensate viscosity continuously increased with the age of the condensates, leading to dynamical arrest. This evolution is well described by a classical model of stretched-exponential form of stress relaxation in soft glassy materials^49–51^:

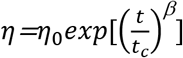

where *η* is the viscosity of the condensates at time *t*, *t_c_* is the characteristic ageing timescale of the condensate, *η*_0_ is the time-independent viscosity, and β is the exponent that quantifies degree of dynamical heterogeneity^51,52^ during condensate ageing. We find that for G3BP1 condensates, β < 0.5 (**Supplementary Table 1**). This behavior is consistent with G3BP1 condensate ageing being governed by a broad distribution of relaxation processes rather than a single characteristic timescale, as previously noted for ageing gels and glasses^53,54^ as well as condensates that are network fluids with low frequency storage and loss moduli that are within an order of magnitude of one another^55,56^. We propose that the continuous increase in viscosity, emergence of sub-diffusive dynamics, and eventual dynamical arrest of G3BP1 condensates is reminiscent of a glass transition ^57^, albeit via ageing rather than cooling.

### G3BP1 condensate ageing is accompanied by an increase in viscous to elastic crossover time

Recently, it has emerged that biomolecular condensates are network fluids^55,56,58,59^. The sequence features of macromolecules dictate the structure and energetics of the interaction network^47^, which gives rise to their emergent material properties^47,56,60,61^. To quantify how network viscoelasticity of G3BP1 condensates changes with ageing, we employed a optical-tweezer-based passive microrheology (pMOT)^60^ assay that we previously developed (**Fig. 2a**). pMOT helps with the extraction of frequency-dependent elastic (Gʹ) and viscous moduli (Gʹʹ) in the frequency range of 0.1 – 100 Hz. This is achieved by acquiring a long-time spatial trajectory and the positional autocorrelation function of the bead optically trapped inside a condensate (**Supplementary Fig. 6**). pMOT measurements revealed that nascent G3BP1 condensates are viscoelastic fluids with Gʹ and Gʹʹ being in the range of 0.10 to 100 Pa within the experimental frequency range. Notably, at frequencies lower than 40 Hz, Gʹʹ dominated over Gʹ, indicating a dominantly viscous long-time, low-frequency behavior of these condensates. Above this frequency, termed as the cross-over frequency (*ω*_crossover_), Gʹ dominates over Gʹʹ (**Fig. 2b**). The inverse of the *ω*_crossover_ defines the terminal relaxation time (*τ*), corresponding to the characteristic time required for the condensate network to dissipate an applied shear stress. For nascent G3BP1 condensates, the characteristic terminal relaxation time *τ* is ∼25 ms.

**Figure 2.**
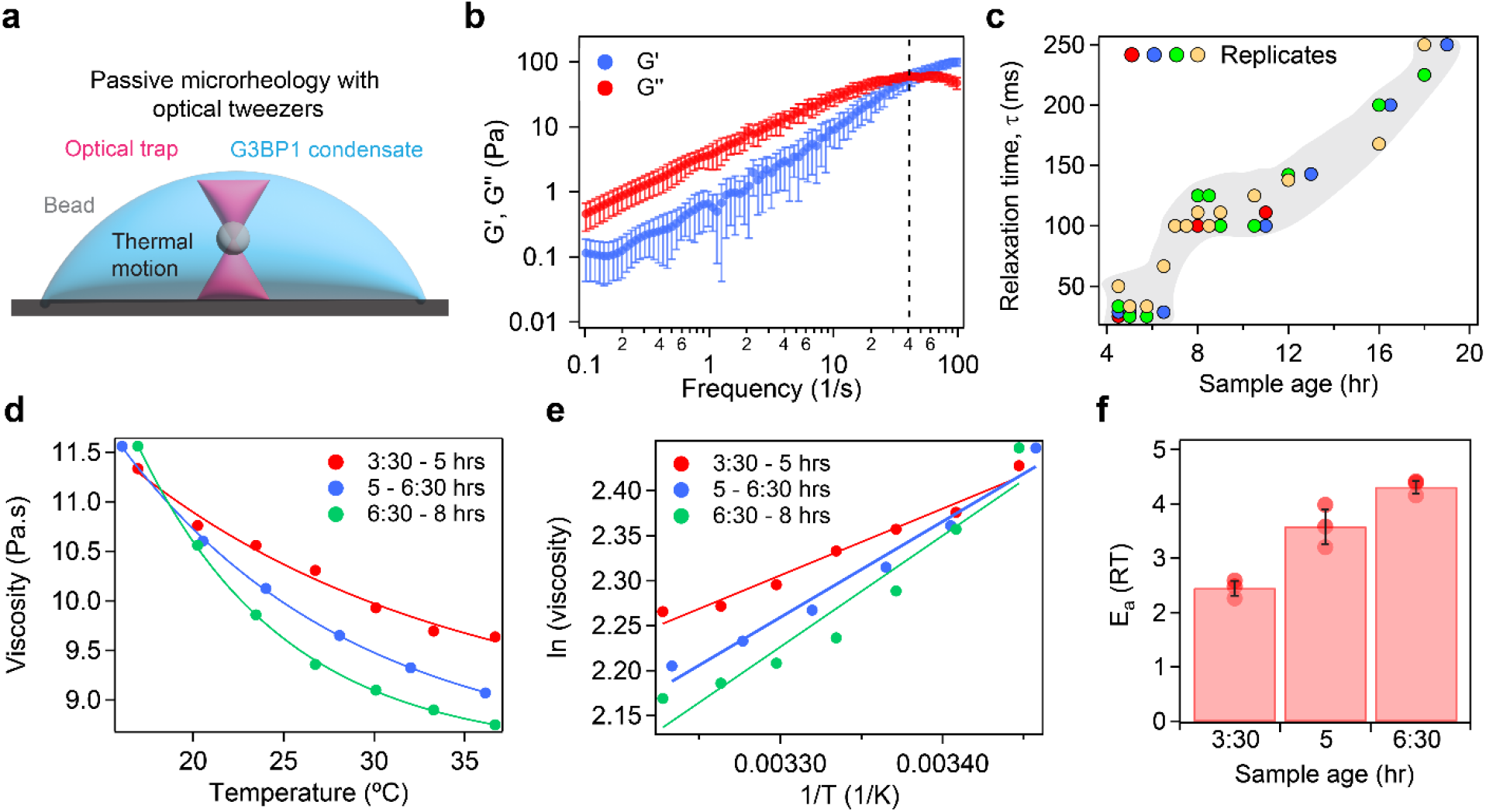
G3BP1 condensate ageing is accompanied by an increase in viscous to elastic crossover time. **a**, Schematic representation of the passive microrheology using optical tweezer (pMOT). **b**, Frequency-dependent storage (Gʹ) and loss (Gʹʹ) modulus of nascent G3BP1 condensates. The points represent the mean values, and the error bars represent the standard deviation (±1 SD) calculated from measurements across nine individual G3BP1 condensates from three independently prepared samples. **c**, The terminal relaxation time (*τ*) of G3BP1 condensates as a function of age. **d**, Viscosity of G3BP1 condensates as a function of temperature, measured using temperature-controlled VPT nanorheology. Points are the data; solid lines are fits to the Arrhenius relation of viscous flow. **e**, Linear fits of *ln*(*viscosity*) vs 1/*T*. The slopes of the lines correspond to the activation energy of network reconfiguration at different sample ages. **f**, Flow activation energy of the ageing condensates at different sample ages. All measurements were repeated using at least three independently prepared samples (*see* Supplementary Figure 8).

Direct measurement of frequency-dependent Gʹ and Gʹʹ for G3BP1 condensates at different sample ages reveal that the terminal relaxation time, *τ* increases by an order of magnitude within 24 hr of condensate age (**Fig. 2c; Supplementary Fig. 7a-f**). Thus, the inter-G3BP1 interaction network relaxes slowly under an applied shear stress with increasing sample age. Notably, G3BP1 condensate ageing obeys time-ageing-time superposition, whereby the Gʹ and Gʹʹ spectra measured at different sample ages collapse onto a single master curve upon horizontal shifting along the frequency axis (**Supplementary Fig. 7g)**^62^. The increment in *τ* with sample age may stem from an increase in the energy barrier that slows the breaking and remaking of physical interactions, which is suggestive of an effective strengthening of the interaction network with condensate ageing. To test for this possibility, we performed temperature-controlled VPT nanorheology and estimated the energy barrier for network reconfiguration^63^, termed flow activation energy (*E_a_*) as a function of ageing (**Fig. 2d, Supplementary Fig. 8a-b**). The viscosity of G3BP1 condensates as a function of temperature follows an Arrhenius-like exponential scaling between 15 and 37 °C. The slope of the ln(*η*) *vs* 1/ *T* curves (**Fig. 2e, Supplementary Fig. 8c-d**) allowed us to estimate the *E_a_*. With condensate age from 0 to 8 hr, *E_a_* increases from ∼ 2.5 to ∼ 4 *RT*, indicating that the energy barrier for the G3BP1 network to reconfigure is ∼ 1.6-fold higher for 8 hr old condensates (**Fig. 2f**). We note that, estimation of *E_a_* beyond these timepoints was not successful because of the rapid ageing of the condensate samples relative to the duration (∼ 1.5 hr for each sample) of each experiment.

### Ageing of G3BP1 condensate is accompanied by the formation of nanoscale cages

We hypothesized that our observations from VPT nanorheology revealing the onset of sub-diffusivity in the bead motion at ∼24 hr might stem from the dynamical growth of nanoscale cages that enable bead confinement in the ageing condensates. To test this hypothesis, we analyzed the bead trajectories at different condensate ages. This revealed a progressive shrinking in the probability distribution of the Brownian fluctuations (**Fig. 3a**). Similar behaviors were also evident from the normalized probability distribution of the beads at different ageing timepoints (**Fig. 3b**). The MSDs corresponding to the probability distribution at the end of 100 s trajectories show a monotonic decrease with sample age (**Fig. 3c**). We noted that the MSDs of the probe particles at 33:40 hr, a timepoint at which they were dynamically arrested, is ∼ 20 nm (**Fig. 3c; inset**). These results indicate that the ageing of G3BP1 condensates may be accompanied by the presence of nanoscale cages that confine the motion of the probe particles.

**Figure 3.**
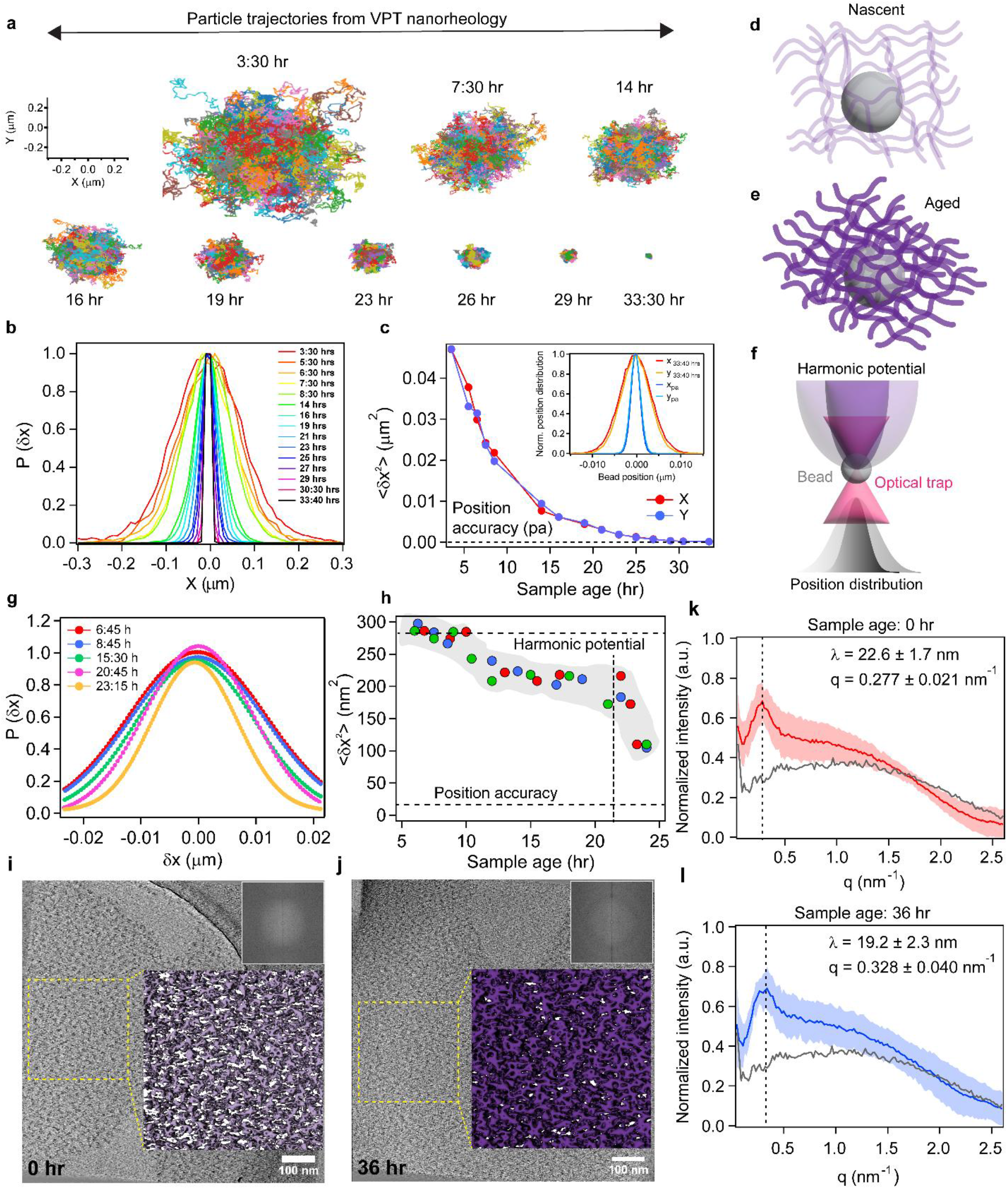
Ageing of G3BP1 condensate is accompanied by the formation of nanoscale cages. **a**, Bead trajectories from VPT nanorheology at different sample ages depict a growing confinement in the condensate microenvironment. **b**, Normalized probability distribution of the beads at different sample ages. **c**, Mean-squared displacements (MSDs) of the beads plotted as a function of sample age. **d-e**, Schematic representation of increasing confinement of bead motion in nascent (d) and aged (e) G3BP1 condensate network. The light and dark colored condensate network in **d** and **e** represent the extent of ageing of the network. **f**, Schematic representation of a passive microrheology using optical tweezer assay showing a single bead in an optical trap inside a G3BP1 condensate. The light and dark purple shadows depict the extent of bead motion inside an optical trap without and with confinement, respectively. The Gaussian distributions in light and dark show the normalized probability distributions of the bead without and with confinement, respectively. **g**, Normalized probability distribution of bead positions in the optical trap at different condensate ages up to the onset of sub-diffusive behavior. **h**, The asymptotic MSDs of the bead in the optical trap at different condensate ages. The upper horizontal dashed lines represent the asymptotic MSD limit set by the harmonic potential of the optical trap, the lower horizontal dashed line corresponds to the position accuracy of the optical trap, and the vertical dashed line represents the sample age corresponding to the emergence of nanocages. **i**, Central slice from a cryo-electron tomogram of freshly prepared G3BP1 condensates imaged within a Cryo-EM grid. The inset shows the 3D tomogram image of the yellow-dashed region. The inset shows the enlarged region, highlighting nanoscale structural arrangements. **j**, Central slice from a cryo-electron tomogram of aged (36 h) G3BP1 condensates, with an inset showing the 3D tomogram image of the yellow-dashed region. **k**, azimuthally averaged radial intensity profile from the two-dimensional FFT of fresh condensates (n = 5). The dominant feature corresponds to a characteristic network spacing of λ = 22.6 ± 1.7 nm. **l**, azimuthally averaged radial intensity profile from aged condensates (n = 5). The characteristic network spacing shifts to λ = 19.2 ± 2.3 nm. All measurements were repeated using at least three independently prepared samples.

The apparent arrest of passive probe particles could arise from progressively slower dissipative stress relaxation, confinement within transient cages formed by the ageing G3BP1 network, or a combination of both^64^. To test this directly, we designed an assay to monitor the asymptotic trajectories of a bead trapped in a harmonic potential of an optical trap inside a G3BP1 condensate^64^. Unlike VPT nanorheology, the probe particle in this case is subjected to the harmonic force exerted by the optical trap, and as such, the stiffness of the optical trap controls the motion of the particle (**Fig. 3d**) in the absence of an equivalent elastic force. In the asymptotic limit, a particle undergoing Brownian motion in a dominantly viscous fluid is expected to be confined by the width of the harmonic potential set by the optical trap (**Fig. 3e**). However, in the presence of a dominantly elastic confinement in the aged G3BP1 condensate, the probe particle will be subjected to an additional harmonic force set by the viscoelastic network. Under this condition, the asymptotic motion of the probe particle will be dominated by the stronger harmonic force (**Fig. 3f**). If the elastic force of the aged condensate dominates over the optical force, the bead motion will register a growing confinement smaller than what is set by the harmonic potential of the optical trap. We analyzed the bead trajectories extracted from pMOT measurements, which show Gaussian fluctuations at all ages. We observed a prominent decrease in the normalized probability distribution of the bead motion at condensate ages at which sub-diffusivity was recorded in VPT nanorheology (**Fig. 3g**). This quantification of the asymptotic MSD as a function of the sample age shows growing elastic confinement of the probe particles. From these measurements, we estimate the dimension of the nanocages on the order of 10-15 nm (**Fig. 3h**). We note that this estimated cage size does not correspond to a physical pore size within the condensate. Rather, it represents the localization length of the probe particle arising from the collective elastic restoring forces exerted by the surrounding G3BP1 network. Although the probe particle is 200 nm in diameter for VPT and 1 µm in the case of pMOT, its Brownian motion is limited to nanometer-scale fluctuations about an equilibrium position, analogous to caging in colloidal glasses.

To visualize the internal organization of G3BP1 condensate network as a function of ageing, we employed cryogenic electron tomography (cryo-ET), which enables three-dimensional imaging of ultrastructure within intact biomolecular condensates^65^. We found that G3BP1 condensates comprise a heterogeneous internal network whose architecture evolves with age. (**Fig. 3i**; lower magnification cryo-ET micrographs at different sample ages are shown in **Supplementary Fig. 9**). Enlarged cryo-ET tomograms reveal a continuous protein network throughout the interior of G3BP1 condensates with visible void spaces (**Fig. 3i, inset)**^31^. Aged condensates collected after 36 hr exhibit a similar overall appearance in tomographic slices (**Fig. 3j**), and enlarged regions likewise show preserved ultrastructural network (**Fig. 3j, inset**). To quantify these differences, we performed azimuthal averaging of the two-dimensional FFTs from tomogram slices, allowing for an unbiased measurement of the characteristic length scale associated with the internal network architecture^66^. The nascent condensates display a dominant peak corresponding to a characteristic spacing of 22.6 ± 1.7 nm (**Fig. 3k**), consistent with a more open network. In contrast, aged condensates exhibit a clear shift to a shorter characteristic length scale of 19.2 ± 2.3 nm (**Fig. 3l**), reflecting nanoscale compaction of the network while maintaining the similar topological features of the dense phase network. Together with our rheology results, these analyses demonstrate that G3BP1 condensates preserve their overall network structure as they age into a dynamically arrested state accompanied by the formation of nanoscale cages. We propose that the initial formation of the condensate network establishes the dimensions of the nanoscale cages, whereas subsequent ageing progressively strengthens the network, giving rise to elastic memory.

### Active microrheology (aMOT) reveals age-dependent emergence of elastic memory in G3BP1 condensate network

The emergence of nanoscale cages, elastic confinement, and slowed network relaxation during condensate ageing suggests the progressive buildup of elastic memory in G3BP1 condensates. However, passive measurements cannot directly assess this dynamical response. To directly probe the age-dependent emergence of elastic memory, we employed active microrheology (aMOT) using an optical-tweezer–based creep test^67^ (**Fig. 4a**). In this assay, a polystyrene bead embedded within the condensate is optically trapped and translated through the condensate at a constant speed. The displacement of the particle relative to the moving trap is monitored while simultaneously recording the forces experienced by the probe particle. In a viscous liquid, the bead follows the trap and remains at its displaced position once the trap motion is stopped. This is true as long as the assay operates within the stable trapping regime where the Stokes drag does not exceed the trap restoring force (**Supplementary Fig. 10)**. For a viscoelastic fluid in the viscous regime, the bead follows the trap with a lag determined by the network relaxation time. In contrast, when the network relaxation time exceeds the experimental timescale, the condensate responds predominantly elastically, and the bead initially follows the moving trap but only over a limited displacement. Once the elastic restoring forces generated by the network exceed the harmonic restoring force of the trap, the bead is lost from the trap, and it recoils toward its original position (**Supplementary Fig. 10)**. Materials with intermediate viscoelastic properties are expected to exhibit a combination of these behaviors.

**Figure 4.**
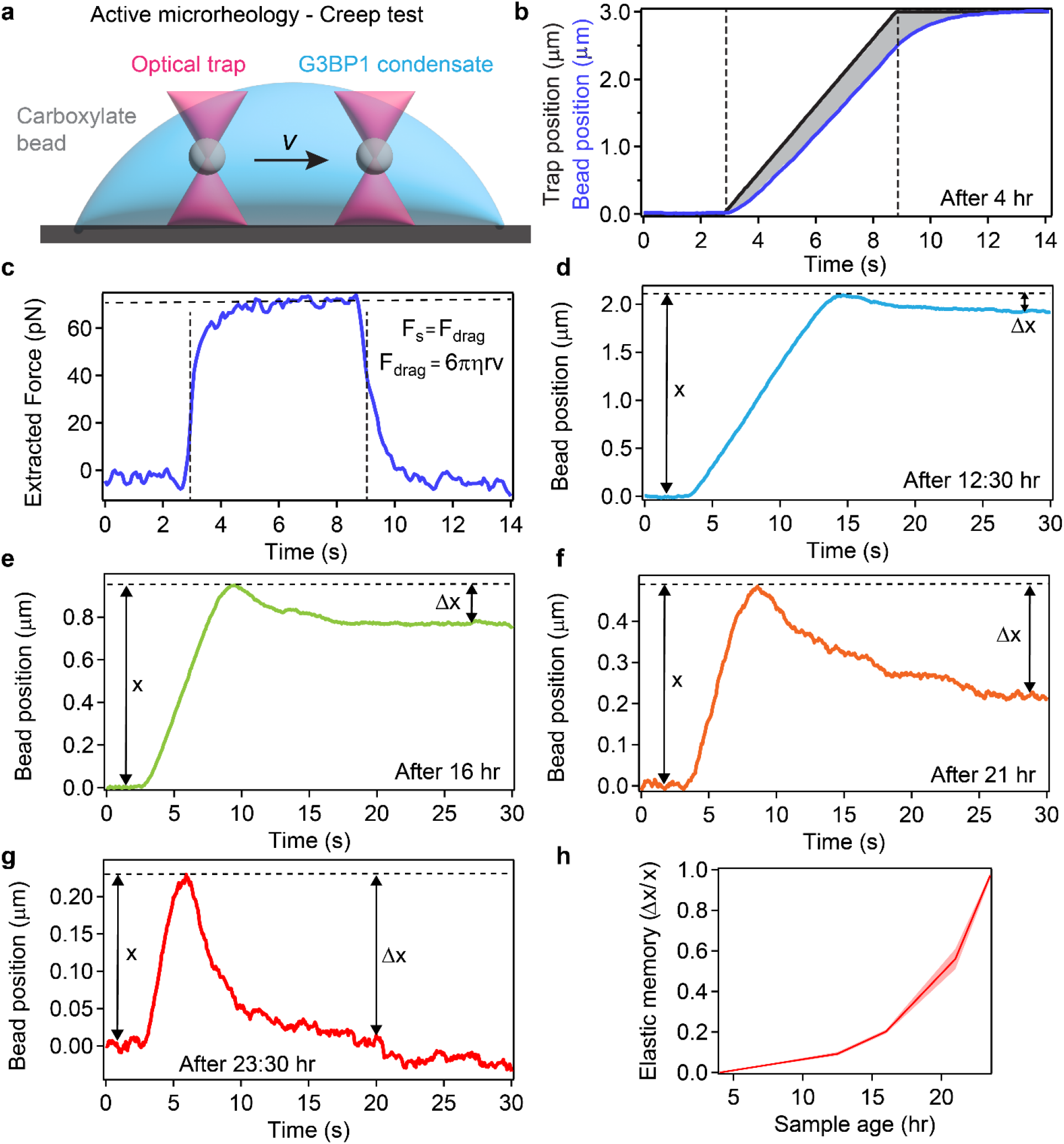
Active microrheology (aMOT) reveals age-dependent emergence of elastic memory in G3BP1 condensate network. **a**, Schematic representation of optical tweezer–based creep test. **b**, Real-time position of the optical trap during a creep test where a 1 µm bead is trapped and moved to 3 µm inside a 4 h old G3BP1 condensate with a constant velocity of 0.5 µm/s (black line). The start and the end time of the trap motion is indicated by vertical dashed lines. Blue line shows the real-time position of the trapped bead. The bead follows the trap with a lag time, which is indicated by the grey shade. **c**, The measured force experienced by the bead during the creep test. The plateau force is used to estimate condensate viscosity, which confirms linear viscoelastic regime. **d-g**, Real-time position of the trapped bead after **d**, 12:30 hrs. **e**, 16 hrs. **f**, 21 hrs. and **g**, 23:30 hrs. of the sample preparation. In these profiles, *x* represents the bead position relative to the starting point before it is lost from the trap, and *Δx* represents the recoiling distance after the bead is released from the trap. **h**, An elastic memory parameter of the bead estimated at different ageing timepoints from measurements shown in **b, d-g**. All measurements were repeated using at least three independently prepared samples.

To test material response of G3BP1 condensates as a function of age, we first programmed the trap to move a 1 µm bead inside a 4 hr old G3BP1 condensate to a maximum distance of 3 µm at a constant speed of 0.5 µm/s (**Fig. 4b**). We observe that the bead follows the trap with a lag, a behavior that is consistent with nascent G3BP1 condensates being viscoelastic fluids (**Fig. 2b**) When operating in the linear viscoelastic regime, the Stokes drag force experienced by the bead is expected to directly yield condensate viscosity. We estimated the viscosity of 4 hr old G3BP1 condensates using aMOT, and this was found to be 7.04 ± 0.58 Pa.s (**Fig. 4c**), which is consistent with the viscosity estimated independently with the VPT nanorheology and pMOT measurements (**Figs**. **1** and **2**).

The creep test results for G3BP1 condensates at increasing age showed markedly different behaviors. At 12.5 hr, the bead failed to reach the programmed displacement of 3 µm (**Fig. 4d**) and exhibited a small recoil after it was released from the trap, indicating the emergence of elastic memory as the condensate network begins to age. This phenomenon is more pronounced with the age of the sample (**Fig. 4e-4f**), where the bead is lost from the trap at progressively smaller distances, followed by an enhanced recoiling of the bead. At 24 hr, when we observed the onset of nanocage formation (**Fig. 3**), we also observed that the bead was only able to move ∼ 200 nm and recoiled back completely to its initial position after trap release (**Fig. 4g**). Collectively, these results allowed direct visualization and quantification of the growing elastic memory (**Fig. 4h**) of G3BP1 condensate network that accompanied their ageing to a dynamically arrested state. Importantly, varying the trap velocity at a fixed sample age revealed that bead recoil decreased progressively as the deformation rate was reduced (**Supplementary Fig. 10g–i**), demonstrating that elastic memory is controlled by the Deborah number (*D_e_* = τ/t), with recoil becoming progressively weaker as the timescale of deformation (t) exceeds the network relaxation time (τ).

### Interaction-domain specific buffering of heterotypic G3BP1 condensate ageing

Recent studies have shown that proteins containing RRMs and IDRs can exhibit microphase separation, giving rise to size-limited nanoscale domains^68,69^. This behavior has been attributed to sequence-and protein architecture-specific inter-domain interactions that feature inter-domain attractions and repulsions^68^. Microphase separation has also been proposed to emerge in polymer networks as a dynamical transition in which elastic forces balance the intrinsic driving forces for phase separation^70^. Motivated by these observations, we used atomistic simulations to determine the pattern of inter-domain interactions in G3BP1 (**see Methods section for details**). The computed pairwise inter-domain interaction coefficients (Δ_XY_ between domains X and Y) are such that -1 ≤ Δ_XY_ ≤ +1. Negative values imply inter-domain attractions, whereas positive values imply repulsions. Values of Δ_XY_ close to zero or with a magnitude of ≈ 0.1 refer to nearly ideal interactions. G3BP1 is a penta-block copolymer, and the sequence-and architecture-specific interactions combine weak and strong inter-domain attractions and repulsions (**Fig. 5a**). The strongest inter-domain attractions are heterotypic, and they mainly involve the NTF2L domain as well as Arg-rich IDR3. All but one of the inter-domain homotypic interactions are repulsive, and the strongest repulsions involve the acidic IDR1. These competing attractive and repulsive interactions are expected to promote dynamical heterogeneity within the ageing condensate network. In polymer networks, this behavior has been linked to microphase-like organization arising from the interplay between elastic constraints and thermodynamic driving forces.

**Figure 5.**
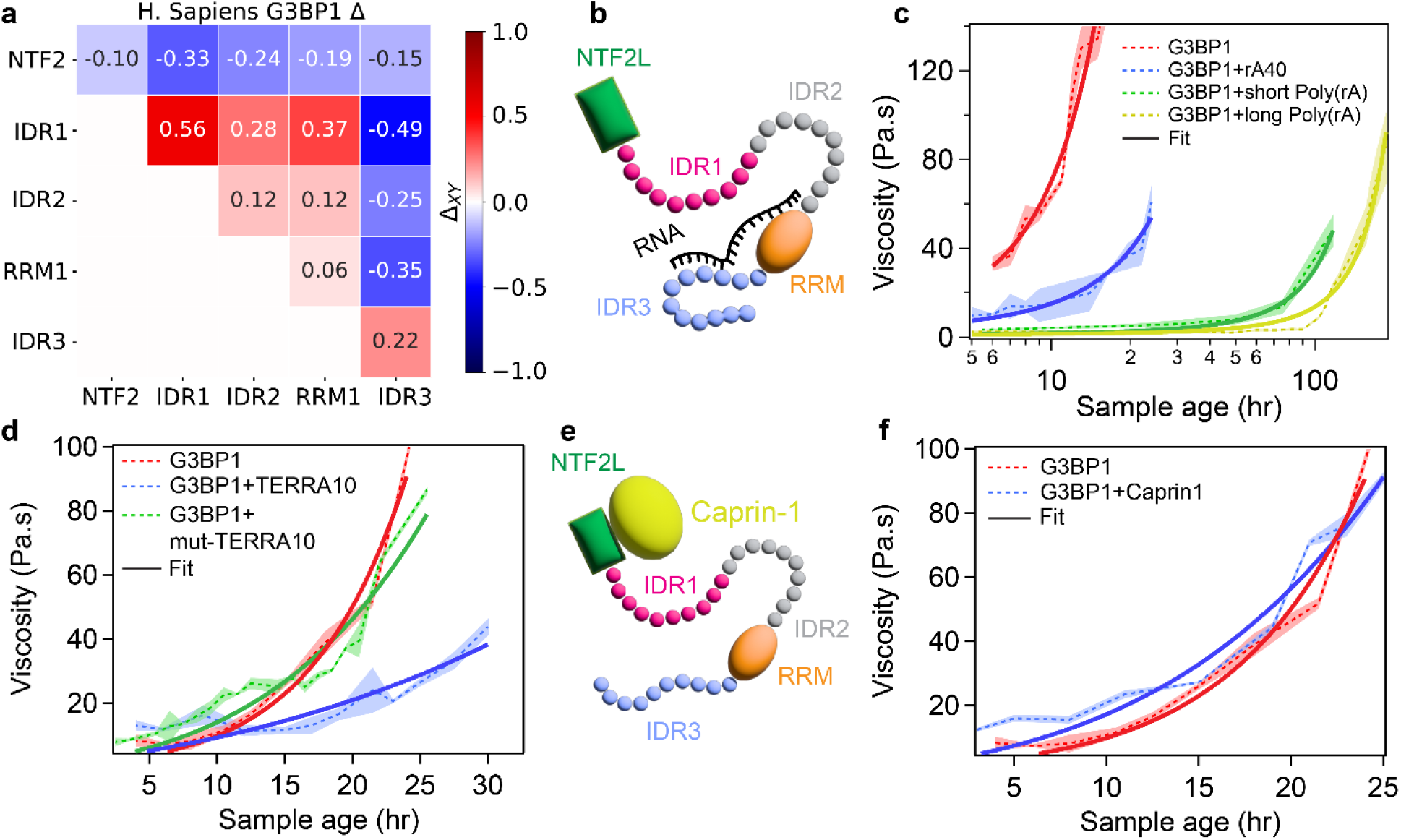
Interaction domain-specific buffering of heterotypic G3BP1 condensate ageing. **a**, Pairwise inter-domain interaction matrix of G3BP1 computed using atomistic simulations. **b,** Schematic representation of G3BP1 interaction domains and an RNA molecule interacting with the RRM and R/G-rich IDR of G3BP1. **c**, Comparison of the viscosity change of G3BP1-RNA condensates with sample age for poly(rA) of three different lengths. **d**, Comparison of the viscosity increase of G3BP1-RNA condensates with sample age for a G-quadruplex-forming RNA (TERRA) and a mutant (mut-TERRA) that lacks the ability to adopt a stable G-quadruplex structure. **e**, Schematic representation of interaction of Caprin-1 with the NTF2L dimerization domain of G3BP1. **f**, Comparison of the viscosity change of G3BP1-Caprin 1 condensates with sample age with G3BP1 condensates alone. In panels c, d, and f, the dotted lines represent the mean viscosity, and the solid line represents the fit using the equation *η =* 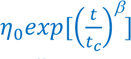. All the measurements were repeated over at least three independently prepared sample replicates.

A plausible model is one where crowder-induced depletion-mediated interactions drive G3BP1 condensation, and the attractive intermolecular, inter-domain interactions, when they form despite being opposed by repulsive interactions, create dynamically organized microphases or clusters within condensate interiors. In support of this model, it is noteworthy that IDR1 is enriched in acidic residues, whereas RRM1 has a positively charged surface and IDR3 is basic, suggesting that electrostatic attractions enable the clustering of G3BP1 molecules whereas repulsions inhibit this clustering. We therefore hypothesize that G3BP1 condensate ageing is governed by barriers that emerge from the competing electrostatic attractions and repulsions within the condensate. To test this idea, we examined the ageing dynamics of G3BP1 condensates under reduced ionic strength (50 mM NaCl) and compared them with those formed at physiological salt concentration (150 mM NaCl). We observed that condensate ageing is accelerated at 50 mM NaCl, with the onset of dynamical arrest occurring at ∼12 hr timescale (**Supplementary Fig. 11**). This behavior is consistent with enhanced electrostatic interactions at lower ionic strength, whereas higher NaCl concentrations weaken these interactions. Because IDR3 is arginine-rich and appears to play a dominant role in G3BP1 homotypic interactions, we further monitored the ageing of condensates in the presence of 5 mM and 10 mM L-arginine (L-Arg), which provides a competing interaction in *trans*, similar to the microphases formed by SRSF1^68^, another RNA-binding protein. In these experiments, L-Arg delayed condensate dynamical arrest in a dose-dependent manner (**Supplementary Fig. 12)**, supporting our hypothesis for the IDR3-driven electrostatic interactions that drive G3BP1 condensate ageing.

G3BP1 is known to interact with RNA, albeit with broad specificity through the RRM and the R/G-rich IDR3 domain^32^ (**Fig. 5b**). We tested the effect of different RNA-binding partners on G3BP1 condensate ageing. We first chose poly(rA) RNAs of different lengths to mimic the interaction between G3BP1 with poly(rA) tail of mRNAs in SGs. In the presence of 200 ng/µL RNA, corresponding to 13.5:1 mass ratio of protein-to-RNA (chosen based on the previous reports on conditions that promote G3BP1-RNA co-condensation^32^), heterotypic G3BP1 condensates remain similar in size as compared to G3BP1 condensates lacking RNA. We also confirmed that Ficoll-70 is largely excluded from G3BP1-RNA co-condensates (**Supplementary Fig. 13**). Employing VPT nanorheology, we observed that poly(rA) RNA significantly slows the ageing of G3BP1 condensates. In the presence of 150 mM NaCl, no discernible changes in condensate viscosity were observed even after 5 days of condensate age (**Supplementary Fig. 14**). In the presence of 50 mM NaCl, the timescale for the onset of dynamical arrest shifted to ∼120 hr compared to ∼12 hr in the absence of RNA (**Fig. 5c**). Notably, this chaperoning effect of RNA is length-dependent: a shorter poly(rA) RNA, rA_40_, showed a weaker effect than longer poly(rA) RNA at a fixed protein-to-RNA mass ratio (**Fig. 5c**). We also observed that the ageing timescale of G3BP1-RNA condensates is inversely correlated with the viscosity of the nascent condensates, with lower viscosity corresponding to longer ageing timescale of G3BP1-RNA condensates (**Supplementary Fig. 15**).

In addition to the buffering effects conferred by RNA length, we asked whether RNA structure plays a role in modulating G3BP1 condensate ageing. RNA G-quadruplexes are found in untranslated regions of mRNA^71^, and they are thought to play an important role in regulating translational activities under stress^72^. G-quadruplex RNAs are known to bind preferentially to R/G-rich IDRs^5,73–76^. To test how G-quadruplex RNAs may impact G3BP1 condensate ageing, we used telomeric repeat containing RNA, TERRA, [UUAGGG]_10_ (TERRA10)^77,78^ as a model system^79–81^. Quantitative rheology measurements revealed that TERRA10 is effective in shifting the timescale of the condensate dynamical arrest to ∼52 hr (**Fig. 5d**) compared to ∼36 hr observed in the absence of any RNA. This buffering effect of TERRA10 is dependent on its structure as a mutant version of TERRA10 (mut-TERRA10: [UUAGUG]_10_), which does not form G-quadruplexes^82,83^, did not show any buffering capacity (**Fig. 5d**). Together, these data suggest that RNA can modulate the ageing dynamics of heterotypic G3BP1-RNA condensates in a length-dependent and structure-specific manner.

A pertinent question at this point is whether the slower ageing of heterotypic G3BP1 condensates is an inherent property of multi-component systems^84^. We tested this possibility utilizing protein binding partners of G3BP1. Proteins such as Caprin-1 specifically bind to the NTF2L domain of G3BP1^39^ (**Fig. 5e**), allowing us to use Caprin-1 as a probe to understand how the interactions of a binding partner to the NTF2L domain of G3BP1 affects the ageing of G3BP1 condensates. In the presence of Caprin-1, we observed that the viscosity of nascent condensates is within ∼2-fold of the G3BP1 condensates lacking Caprin-1. Caprin-1 binding to NTF2L domain of G3BP1 enables the recruitment of translationally silent mRNAs and assists in overcoming the autoinhibited state of G3BP1, effectively driving the condensation of large G3BP1-mRNA complexes^31,32,35,85^. Notably, the condensates in the presence of Caprin-1 did not show a significant change in the ageing timescale (**Fig. 5f**). Collectively, these observations indicate that IDR1, IDR3, and the RRM in G3BP1 are the main drivers of G3BP1 condensates ageing, whereas RNA can slow the ageing dynamics by frustrating the molecular interactions mediated by the RRM and IDR3.

### Coarse-grained simulations reveal how different interactions regulate G3BP1 condensate ageing

To gain mechanistic insights into the domain-specific interactions in the absence and presence of RNA that regulate the ageing dynamics of G3BP1 condensates, we deployed coarse-grained simulations by parameterizing a phenomenological G3BP1 model using the inter-domain interaction coefficients derived from atomistic simulations that use the ABSINTH implicit solvation model and forcefield paradigm^86^. The coarse-grained simulations were based on LaSSI^87^, which is a lattice-based Monte Carlo simulation engine (**see Methods section for details**). Each G3BP1 molecule comprises five different bead types that correspond to each G3BP1 domain (**Fig. 6a, top panel**). The pairwise interaction energies between each bead type are defined by the inter-domain interaction coefficients derived from atomistic simulations (**Fig. 5a**). We probed the phase behaviors in coarse-grained simulations as a function of a single scaling parameter *s*, that helps vary the net balance of inter-domain attractions and repulsions. In this approach, the chemical potential, µ(*s*), is a function of the scaling parameter, *s*. As a result, the balance between attractive and repulsive interactions is tuned without altering the intrinsic hierarchy of inter-domain interactions. We fix the size of the simulation volume and the number of molecules, guarding against finite size artifacts^84^. As *s* increases, we observe clusters that are system-spanning (percolated) for lower scaling factors that transition into beads-on-a-string types of morphology at higher scaling factors (*s* ≥ 1.25) (**Fig. 6a, bottom panel**). We observe a rightward shift in the cluster size distribution *p*(*n*). Here, *p*(*n*) quantifies the probability of observing clusters comprising *n* molecules. For the higher scaling factor (*s* = 1.25), we quantified the cluster size distribution and the spatial organization of different G3BP1 domains within the clusters, and this revealed that RRM and IDR3 are localized to the cores whereas NTF2L/IDR1/IDR2 are localized to the surfaces of clusters (**Fig. 6b-6c**). The surface localization of the “sticky” domains provides sites for association and networking with other molecules and clusters.

**Figure 6.**
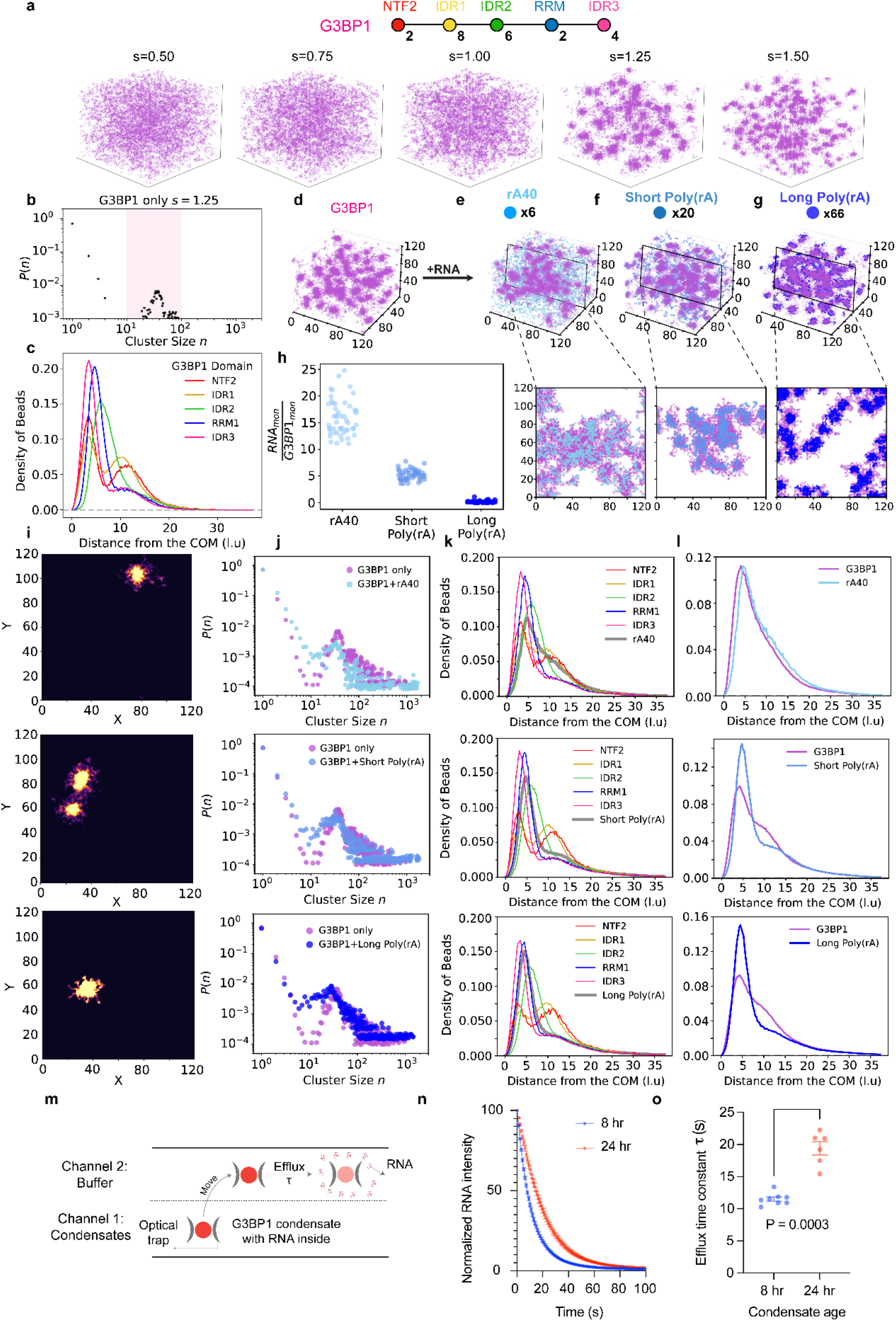
Coarse-grained simulations reveal how different interactions regulate G3BP1 condensate ageing. **a**, Schematic representation of the coarse-grained simulation setup for G3BP1 with the number of beads assigned for the interaction domains as indicated. Also shown are the snapshots of simulation box for different interaction strength between the G3BP1 domains modulated using a scaling factor ‘*s*’ utilizing the inter-domain interaction matrix shown in Figure 5a. **b,** Cluster size distribution for the G3BP1 clusters with *s*=1.25. **c**, Spatial organization of G3BP1 interaction domains quantified in terms of their radial distribution functions. **d,** snapshots of the simulation box corresponding to the G3BP1 at *s*=1.25 and G3BP1-RNA simulations for different RNA lengths **e**, rA40, **f**, short Poly(rA), and **g**, long Poly(rA) with the corresponding insets. **h**, Quantification of the fraction of RNA monomers relative to the fraction of G3BP1 monomers in the cluster-devoid space for three different RNA lengths. **i**, Heat map images of G3BP1-RNA clusters for three different lengths of RNA. **j**, Cluster size distribution of G3BP1-RNA clusters overlayed on the G3BP1-only cluster size distribution for the three RNA lengths. **k**, Radial distribution functions for the G3BP1 interaction domains inside the G3BP1-RNA clusters for the three RNA lengths. **l**, Radial distribution functions of three RNAs overlayed on the radial distribution function corresponding to G3BP1. **m**, Schematic representation of the efflux assay used to quantify the efflux of labeled RNA molecules from G3BP1-RNA condensates as condensates age. **n**, Normalized RNA intensity profiles measured for early age (8 hr after sample preparation) and aged (24 hr after sample preparation) G3BP1-RNA condensates. **o**, Quantification of RNA efflux time constant for early age (8 hr after sample preparation) and aged (24 hr after sample preparation) G3BP1-RNA condensates. The statistical significance between the efflux time constant at 8 hr and 24 hr was determined to be P=0.0003 estimated using Welch’s t-test for N=8 for 8 hr and N=6 for 24 hr conditions. All the measurements were repeated over at least three independently prepared sample replicates.

To investigate the role of RNA in regulating G3BP1–RNA cluster formation, we performed analogous simulations incorporating RNA, in which the chain lengths of RNA-mimicking polymers were selected to match the nucleotide lengths used in the poly(rA) RNA experiments (**Fig. 6d-6g**). This approach allowed us to probe how RNA length modulates the topology of and spatial organization within G3BP1–RNA clusters (**see Methods section for details**). Notably, the only attractive interactions between the RNA beads were with the sites of the RRM1 and IDR3 of G3BP1. All other interactions with G3BP1 beads were set to be zero, which on a lattice become implicitly repulsive (**Supplementary Figure 20**). The simulation snapshots (**Fig. 6d-6g**) provide a visual depiction of G3BP1 alone and G3BP1-RNA heterotypic clusters at different RNA length. The cluster size distributions indicate that RNAs drive the formation of smaller G3BP1 clusters compared to RNA-free conditions (**Fig. 6i-6j**). This effect is most pronounced for the longest RNA mimic (long poly(rA)) at a scaling factor of 1.25. As we vary the scaling factor applied to G3BP1 inter-domain interactions, the presence of RNA promotes the association of G3BP1–RNA clusters (**Supplementary Fig. 16**). Consistent with this behavior, simulation snapshots reveal that RNA molecules localize predominantly within the cluster cores, enriched in regions containing the RRM and IDR3 (**Fig. 6k**). As the length of RNA increases, we observe a higher partitioning of RNA into the G3BP1-RNA clusters, quantified by the number of RNA monomers relative to the number of G3BP1 in the non-cluster phases (**Fig. 6h, l**).

Overall, the simulation results suggest that poly(rA) effectively caps interactions mediated by the RRM and R/G-rich IDR3 of G3BP1 (**Fig. 5b**). A direct consequence of this capping is a reduction in homotypic interactions between G3BP1 molecules within condensates that comprise RNA. If RNA engagement attenuates these homotypic interactions, G3BP1 partitioning into G3BP1–RNA condensates would be expected to decrease relative to G3BP1-only condensates. Indeed, in accord with this hypothesis, we observed ∼2-5 fold reduced G3BP1 partitioning in G3BP1–RNA condensates compared with G3BP1-only condensates (**Supplementary Fig. 17**), with the lowest G3BP1 partitioning observed for longer poly(rA). These results, which show the dilution of G3BP1 in the dense phase^15^, establish a molecular basis for condensate ageing, demonstrating how attractions contribute to heterotypic buffering^84^ via protein-RNA interactions to tune network connectivity, as seen in simulations, and the measured dynamics of ageing.

### Functional implications of G3BP1 condensate dynamical arrest

A direct consequence of SG ageing is an increased resistance to disassembly. These disassembly-resistant states can act as kinetic traps for RBPs and RNAs, with potential implications for disease pathogenesis^89^. Using the G3BP1 condensate system together with a microfluidic-based condensate dissolution and retention assay^25^ (**Fig. 6m**), we tested whether RNA sequestration is altered as condensates age. In this assay, we examined heterotypic G3BP1–poly(rA) condensates at two distinct sample ages (**Fig. 6n**): ∼8 hr, corresponding to a stage at which condensates have not aged significantly, and ∼24 hr, corresponding to the onset of dynamical arrest. Using multichannel microfluidics under laminar flow, we monitored the efflux of poly(rA) from optically-trapped G3BP1 condensates by fluorescence microscopy (**see Methods for assay details**). The condensates remained stable against dissolution under constant flow (**Supplementary Fig. 18**). Notably, at ∼24 hr of sample age, the efflux of poly(rA) from the condensates was ∼2-fold slower than that observed for ∼8 hr-old condensates (**Fig. 6o**). Together, these results indicate that RNA release dynamics are significantly altered as G3BP1 condensates age. The implication is that the release of RNA is slowed by condensate ageing. This highlights the importance of enzymes that actively remodel or degrade RNA as enablers of disassembly of the RNA components of SGs^10,13,90^.

## Conclusion

In this study, we investigated the ageing dynamics of G3BP1 condensates that form via crowder-induced depletion-mediated interactions, and the effects of different heterotypic (RNA vs. protein) interactions on these ageing dynamics. This ageing process is defined by a continuous increase in condensate viscosity and network relaxation times, accompanied by the formation of nanoscale cages and emergence of elastic memory in the condensate network. Cryo-ET shows that dynamically arrested condensates retain a similar network architecture without the emergence of short-or long-range order, implying that ageing primarily prolongs the lifetime of nanoscale cages, consistent with dynamically driven microphase-like organization. We argue that the aged G3BP1 condensates are best described as viscoelastic network glasses^54,91^ because dynamical arrest emerges through progressive slowing of stress relaxation^64^ without an increase in elastic stiffness. While the plateau elastic modulus remains approximately constant (**Supplementary Fig. 6**), the network relaxation time grows markedly with age (**Fig. 2**), leading to persistent elastic memory and recoil in active microrheology measurements. This behavior is consistent with glass-like ageing^52^ of a percolated network, in which progressively slowed relaxation gives rise to dynamically arrested, heterogeneous domains reminiscent of intra-condensate microphase-like organization^2^. Our simulations reveal the potential for nanoscale domains that form via microphase separation and the clustering of these domains with changes to the balance of inter-domain interactions, which appear to be prevalent as probed by the salt-dependence of ageing dynamics.

Previous studies have proposed multiple orthogonal routes for protein condensate ageing. In systems capable of amyloid formation, ageing is often accompanied by fibril formation^27,92,93^. In other cases, condensates have been reported to undergo glass-like transition while maintaining their viscoelastic Maxwell fluid-like character at all ages, exhibiting increased viscosity without changes in elastic modulus^29^. A third route has been described for protein condensates transitioning from a Maxwell fluid to Kelvin-Voigt solid accompanied by a disorder-to-order transition with semi-crystalline structural features^94^. Our observations are broadly consistent with the glass-like ageing described by Jawerth et al.^29^ for PGL-3 condensates. However, the aged G3BP1 condensates exhibit distinctive features, including nanocage formation and the emergence of elastic memory, that extend beyond prior descriptions. The formation and temporal evolution of nanocages in G3BP1 condensates complement recent observations of nanometer-scale domains that regulate molecular diffusion and connectivity within condensates formed by RNA-binding protein FUS^95^. They are also reminiscent of nanoscale hubs that form transiently in condensates formed by prion like low complexity domains^96^, and by the bacterial protein PopZ^97^. Here we observe a dynamical buildup of nanoscale cages that enabled molecular confinement and the buildup of elastic memory during ageing. This appears to be an orthogonal mode for dynamic reorganization of intra-condensate interaction networks that governs their transition toward glass-like dynamical arrest.

From an interaction-network perspective, we propose that electrostatic interactions between the acidic IDR1 and the basic, R/G-rich IDR3 are the primary drivers of inter-G3BP1 interactions that drive ageing. The crowders appear to facilitate homotypic interactions by bringing G3BP1 molecules into proximity via depletion effects^37^, lowering the effective barrier for intermolecular association. Excipients that frustrate the depletion-mediated interactions help slow the ageing process. We showed this by salt titration, using small molecule L-arg, and by systematically introducing binding partners targeting specific G3BP1 interaction domains. We first introduced poly(rA) RNA, which engages the RRM and IDR3, and observed a pronounced delay in G3BP1 condensate ageing, consistent with heterotypic RNA-protein interactions competing with depletion-mediated protein-protein interactions. Notably, RNA length and hence the valence of nucleotides have a clear regulatory role in this context, with longer RNAs providing more effective buffering against ageing. This observed RNA-length dependent chaperoning of G3BP1 condensate ageing carries important biological implications. Recent work suggests that poly(rA) tail extensions are a prerequisite for SG formation^98^. It was shown that acute stress is controlled by a balance between the action of cytoplasmic poly(rA) element binding (CPEB) proteins that elongate these tails and deadenylating (such as the CCR4-NOT and PAN2-3 complexes) that shorten them, resulting in global poly(rA) tail lengthening^99,100^. Our biophysical results directly link poly(rA) tail length to stress-responsive G3BP1 condensate ageing dynamics, providing a functional framework for how RNA processing may modulate SG stability and persistence. In addition to poly(rA) length, we found that G-quadruplex RNAs provide structure-specific chaperoning against condensate ageing. In contrast, Caprin-1, which binds to the NTF2L domain, appears to mildly accelerate ageing by facilitating G3BP1 interactions.

More broadly, our findings suggest that stress-granule persistence is governed not only by the aggregation propensity of recruited client proteins^17,19–21^, but also by the age-dependent remodeling of the interaction network of the scaffold itself. Regulation of G3BP1 interactions by RNA length, binding specificity, and partner proteins may therefore help determine whether stress granules remain dynamic and reversible or progress toward persistent, disease-associated material states.

## Acknowledgements

This work was supported by the US National Institutes of Health (NIH) through grant R35 GM138186 (to P.R.B.), the US National Science Foundation (NSF) through grant MCB-2227268 (to R.V.P.), the St. Jude Children’s Research Collaborative on the Biology and Biophysics of RNP Granules (to P.R.B., R.V.P. and J.P.T.), the Center for Biomolecular Condensates in the McKelvey School of Engineering at Washington University in St. Louis (to R.V.P.), and US National Science Foundation (NSF) through grant ITE-2452698 (to A.R.). We acknowledge the assistance of Dr. Wade Borcherds (SJCRH) regarding the LC-MS experiments of Caprin-1 and methods. We are grateful for critical feedback from the members of the Banerjee, Pappu, Taylor, and Mittag labs. Any opinions, findings, conclusions, or recommendations expressed in this material are those of the author(s) and do not necessarily reflect the views of the NSF, NIH, or any other funding bodies.

## Conflicts of Interest

P.R.B. is a member of the Biophysics Reviews (AIP Publishing) editorial board. This affiliation did not influence the work reported here. All other authors have no conflicts to report.

## Author Contributions

Conceptualization: P.R.B. and A.S.; Methodology – experiments: P.R.B, A.S., A.R., B.G, T.S.M.; Methodology – G3BP1 protein production: H.J.K, B.M.F., and B.G.; Methodology – simulations: R.V.P and V.L. Investigation: P.R.B, A.S., A.R., B.G, T.S.M., R.V.P and V.L.; Data curation: P.R.B, A.S., A.R., T.S.M., R.V.P and V.L.; Formal analysis and critical thinking: P.R.B, A.S., A.R., T.S.M., R.V.P and V.L.; Visualization: P.R.B, A.S., A.R., T.M., R.V.P and V.L.; Writing – original draft: P.R.B. and A.S.; Writing – review & editing: all authors; Supervision: P.R.B. and R.V.P.; Funding acquisition: P.R.B., R.V.P, and J.P.T.

## Data Availability

Unless otherwise stated, all data supporting the results of this study can be found in the article and the supplementary files.

## Code Availability

Codes for nanorheology are available via GitHub at https://github.com/BanerjeeLab-repertoire.

All original code for the simulation analyses can be found on the GitHub repository of the Pappu lab at https://github.com/Pappulab/Competing-molecular-interactions-govern-dynamical-arrest-in-g3bp1-condensates.

## Declaration of Generative AI and AI-assisted technologies

The authors acknowledge the use of Microsoft Copilot and ChatGPT to enhance the readability and clarity of certain sections of the manuscript text. The authors reviewed and edited all content generated with the assistance of AI tools and take full responsibility for the content of this manuscript.

## Notes

### Competing Interest Statement

The authors have declared no competing interest.

