## Supplementary Information for "Competing Molecular Interactions Govern the Dynamical Arrest of G3BP1 Condensates"


#### The PDF file includes:

Materials and Methods
Figs. S1 to S20
Table S1
Captions for Videos S1 to S5
References (1-32)

#### Other Supplementary Materials for this manuscript include the following:

Videos S1 to S5

**(a)**

**G3BP1**

IDR1 - TEPQEESEEEVEEPEERQQTPEVVPDDSGTFYDQAVVSNDME
EHLEEPVAEPEPDPEPEPEQEPVSEIQEEKPEPVLEETAPED

IDR2 - AQKSSSPAPADIAQTVQEDLRTFSWASVTSKNLPPSGAVPVTGI
PPHVVKVPASQPRPESKPESQI PPQRPQRDQRVREQRINIPPQ
RGPRPIREA GEQGDIEPRRMV

IDR3 - EEKKTRAAREGDRRDNRLRGPGGPRGGLGGGMRGPPRGGM
VQKPGFGVGRGLAPRQ

**FUS IDR**

MASNDYTQQATQSYGAYPTQPGQGYSQQSSQPYGQQSYSGYSQSTD
TSGYGQSSYSSYGQSQNTGYGTQSTPQGYGSTGGYGSSQSSQSSYG
QQSSYPGYGQQPAPSSTSGSYGSSSQSSSYGQPQSGSYSQQPSYGG
QQQSYGQQQSYNPPQGYGQQNQYNSSSGGGGGGGGGGNYGQDQS
SMSSGGGSGGGYGNQDQSGGGGSGGYGQQDRG

**hnRNPA1 IDR**

GSMASASSSQRGRSGSGNFGGGRGGGFGGNDNFGRGGNFSGRGGFG
GSRGGGGYGGSGDGYNGFGNDGSNFGGGGSYNDFGNYNNQSSNFGP
MKGGNFGGRSSGPYGGGGQYFAKPRNQGGYGGSSSSSSYGSGRRF

**TDP-43 IDR**

RFGGNPGGFGNQGGFGNSRGGGAGLGNNQGSNMGGGMNFGAF
SINPAMMAAAQAALQSSWGMMGMLASQQNQSGPSGNNQNQGNM
QREPNQAFGSGNNSYSGSNSGAAIGWGSASNAGSGSGFNGGFGSS
MDSKSSGWGM

**(b)**

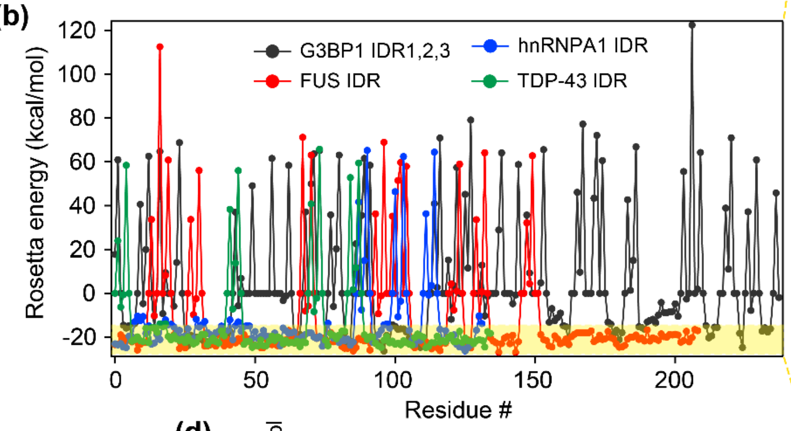


**(c)**

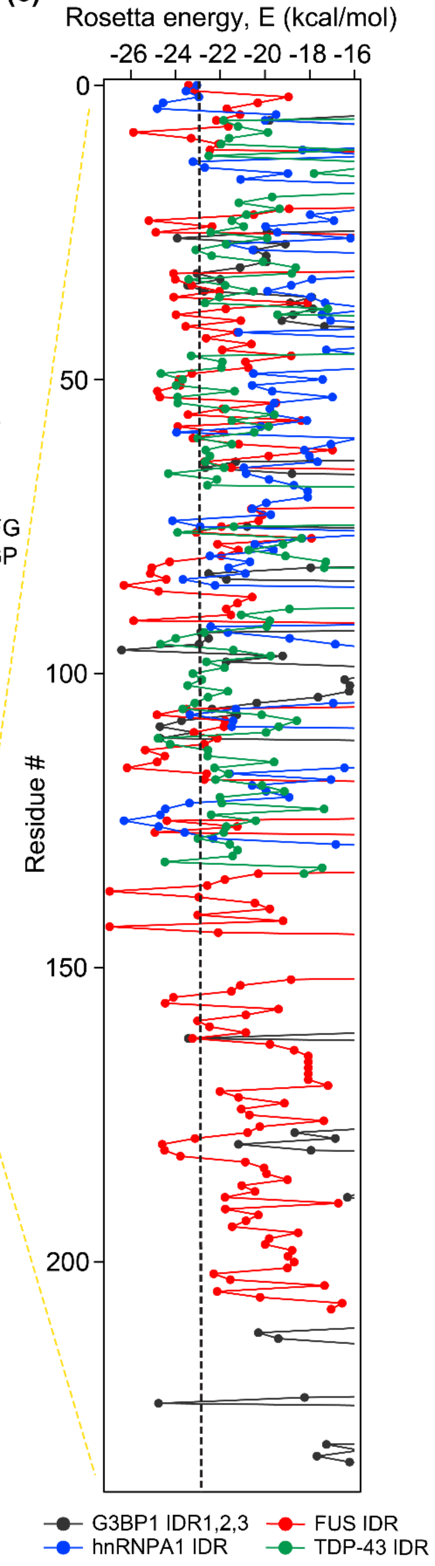


**(d)**

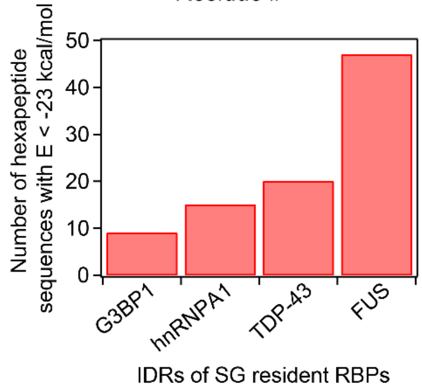

**Supplementary Figure 1. G3BP1 IDRs have a lower number of hexapeptides favoring zipper formation than other RBPs prone to form fibrils.** (a) Sequences of IDRs of G3BP1, FUS, hnRNPA1, and TDP-43, which are SG resident RBPs. (b) Rosetta energy (kcal/mol) profiles for the IDR sequences of the four RBPs determined from ZipperDB[1] (https://zipperdb.mbi.ucla.edu/). (c) Zoomed-in view of the Rosetta energy profile to illustrate the number of hexapeptides with Rosetta energy < -23 kcal/mol, which is the threshold for stronger zipper formation propensity by a hexapeptide. The dotted line corresponds to this threshold of -23 kcal/mol. (d) Number of hexapeptide sequences with Rosetta energy < -23 kcal/mol in the IDR sequences of the four RBPs.

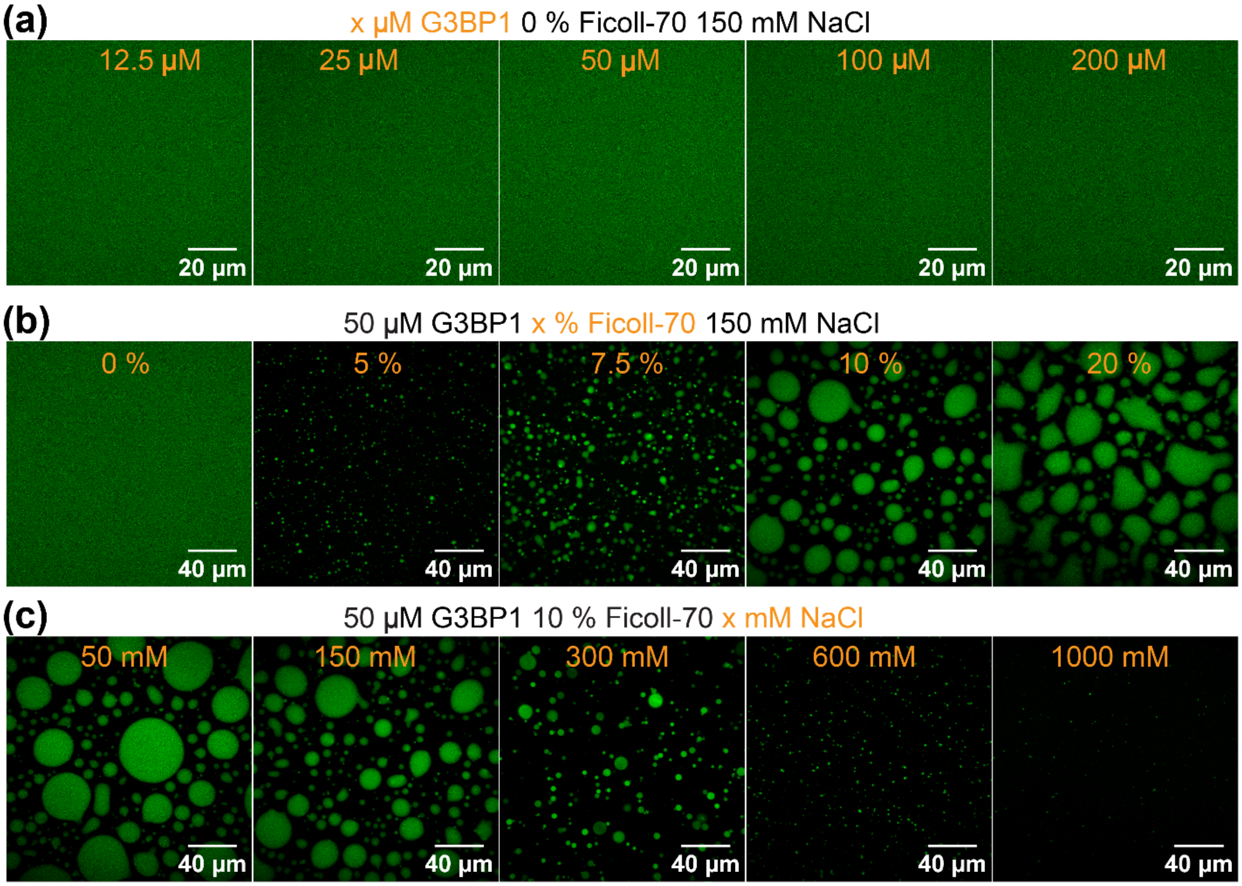


**Supplementary Figure 2. Concentration, crowder, and salt-dependent phase separation of G3BP1.** (a) Phase behavior of G3BP1 as a function of protein concentration at 150 mM NaCl and in the absence of crowder (0 % Ficoll-70). (b) Phase behavior of G3BP1 as a function of Ficoll-70 concentration at 50 μM G3BP1 and 150 mM NaCl. (c) Phase behavior of G3BP1 as a function of NaCl concentration at 50 μM G3BP1 and 10 % Ficoll-70.

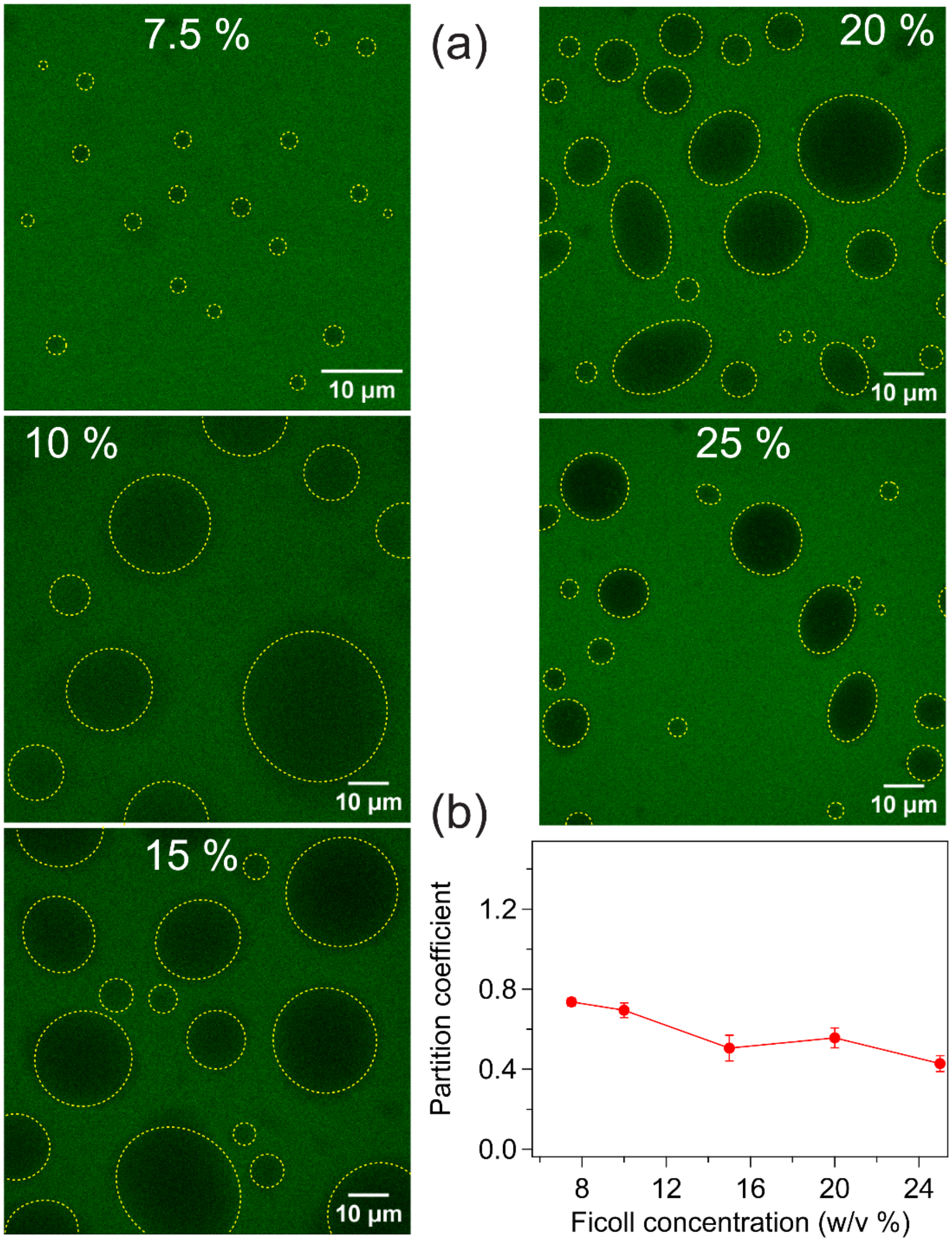


**Supplementary Figure 3. Ficoll-70 promotes the formation of homotypic G3BP1 condensates**. (a) Partitioning of Fluorescein-labeled Ficoll-70 inside G3BP1 condensates at different concentrations of Ficoll-70, images shown for 7.5%, 10%, 15%, 20%, and 25% (w/v) Ficoll-70 concentration. (b) Partition coefficient of Ficoll-70 inside G3BP1 condensates as a function of Ficoll-70 concentration.

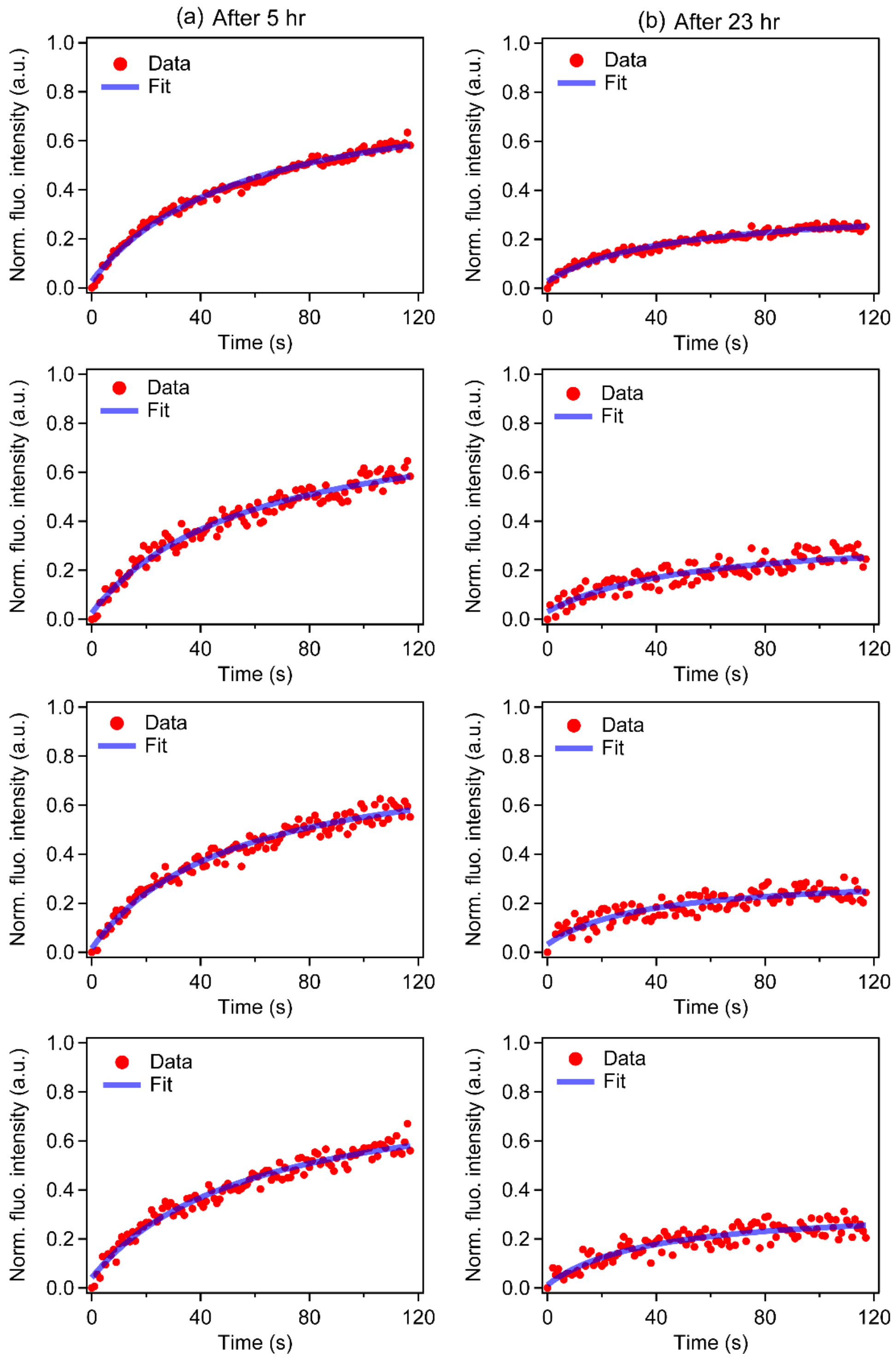


**Supplementary Figure 4. FRAP recovery curves for G3BP1 condensates at different sample ages.** Individual fluorescence recovery curves and their fit using the equation $I_{recovery}(t) = \frac{I_0 + I_\infty \frac{t}{\tau_{1/2}}}{1 + \frac{t}{\tau_{\frac{1}{2}}}}$ (see Methods section) after photobleaching Atto488-labeled G3BP1 inside homotypic G3BP1 condensates after (a) 5 hr and (b) 23 hr of sample preparation.

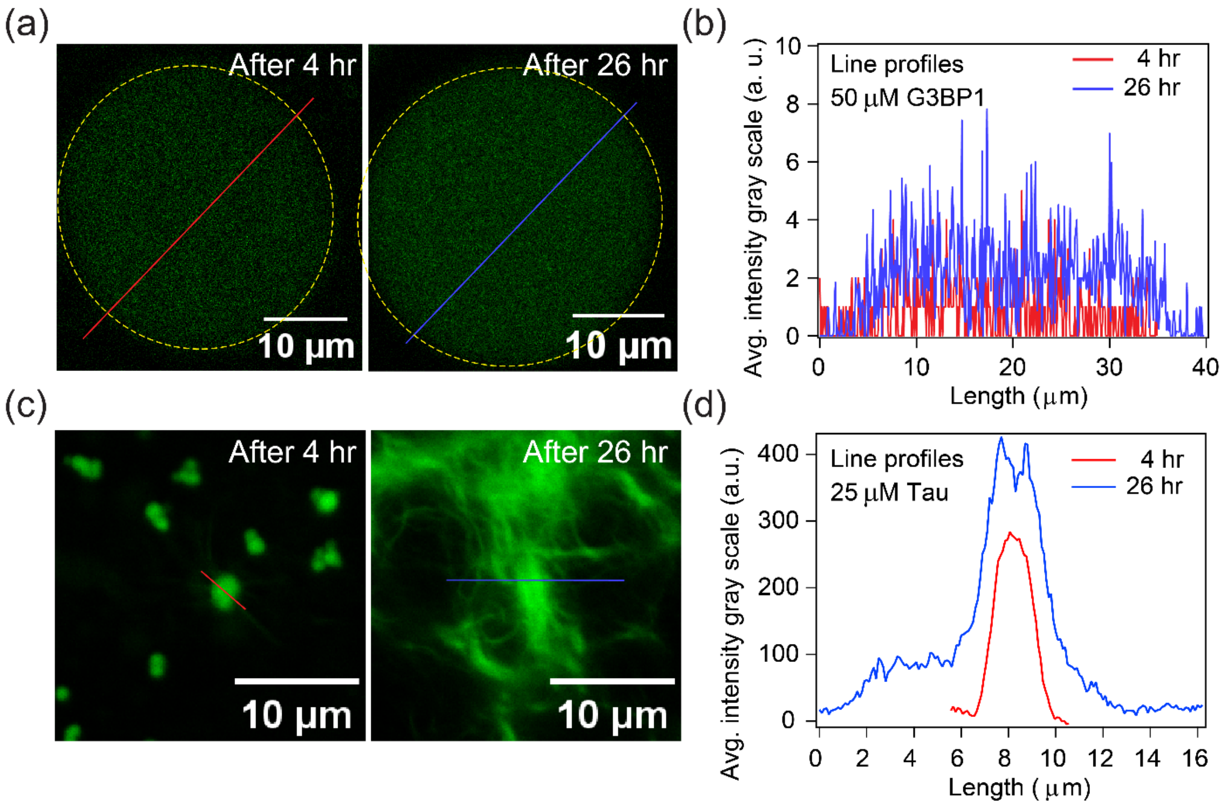


**Supplementary Figure 5. Aged G3BP1 condensates are not ThT-positive.** (a) Thioflavin-T (ThT) fluorescence intensity from 50 μM G3BP1 condensates after 4 hr and 26 hr of sample age. (b) Intensity profiles corresponding to the lines through the condensates as shown in (a). (c) ThT fluorescence intensity from 25 μM Tau condensates, used as a positive control, after 4 hr and 26 hr of sample preparation. (d) Intensity profiles corresponding to the line through the condensates after 4 hr and fibrils after 26 hr as shown in (c). All ThT images were collected utilizing identical experimental conditions and image acquisition settings.

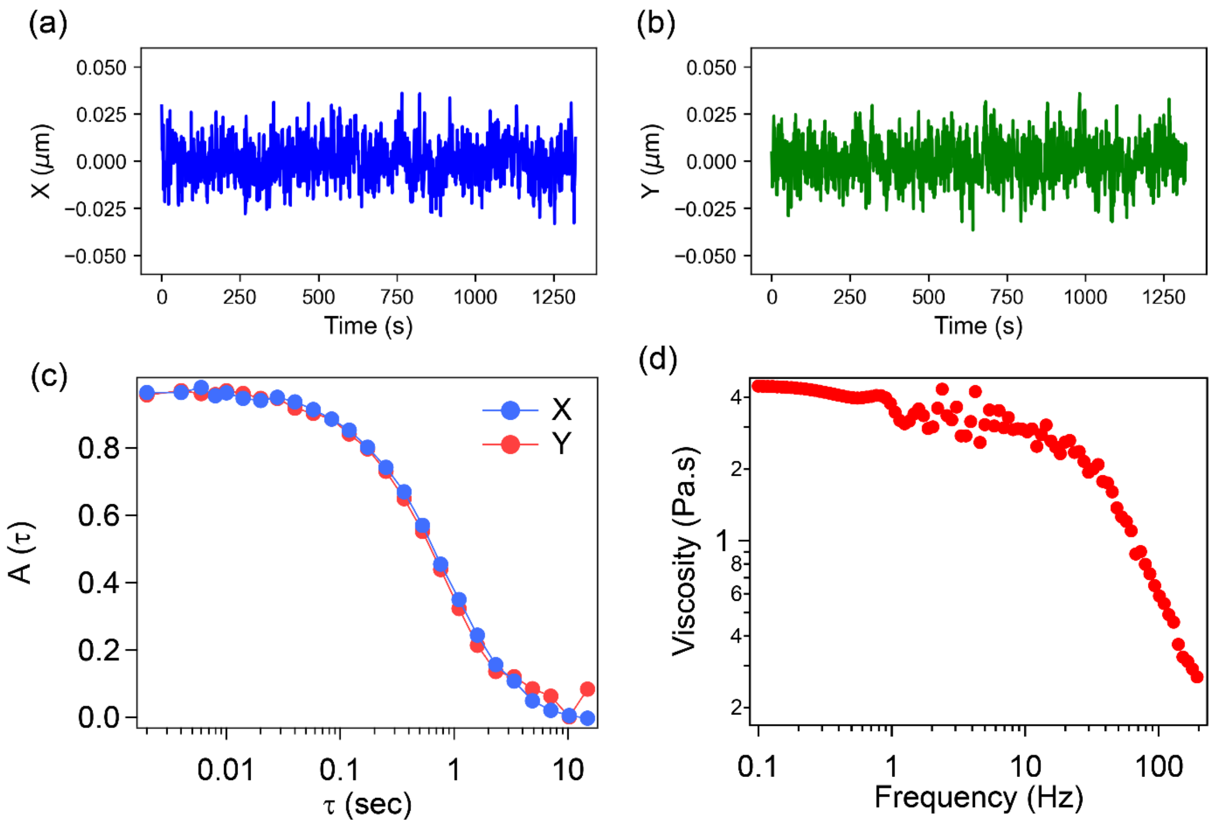


**Supplementary Figure 6**. **Representative pMOT data for viscoelastic characterization of G3BP1 condensates**. Trajectory of a 1 μm bead embedded inside G3BP1 condensate in (a) X and (b) Y. (c) Autocorrelation curve of the bead in X and Y estimated from the bead trajectories shown in (a) and (b). (d) Frequency-dependent viscosity of the G3BP1 condensates determined from pMOT measurements.

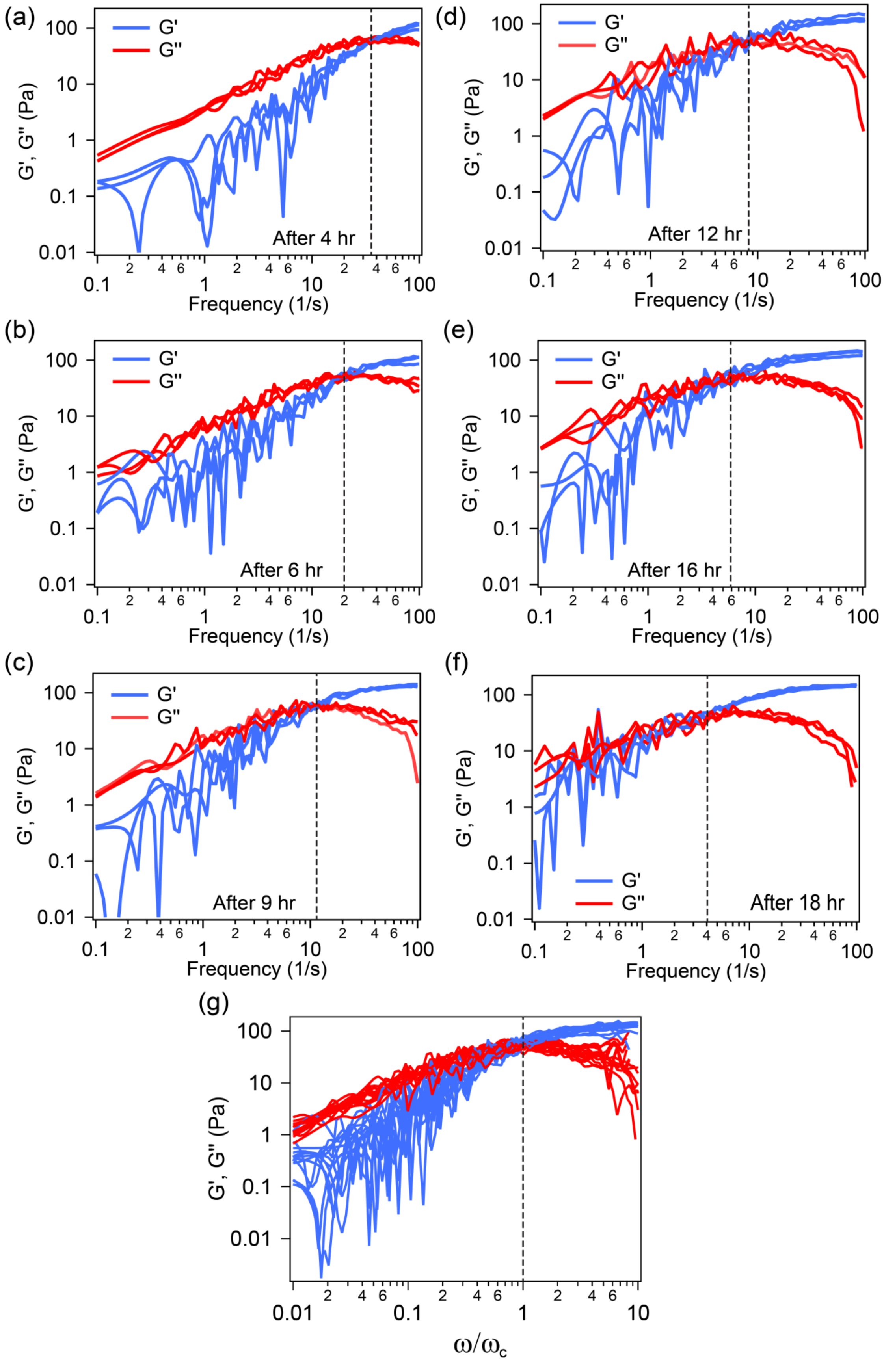

(a)
G'
G''
After 4 hr
G', G'' (Pa)
Frequency (1/s)
(b)
After 6 hr
(c)
After 9 hr
(d)
After 12 hr
(e)
After 16 hr
(f)
After 18 hr
(g)
ω/ω_c

**Supplementary Figure 7. Frequency-dependent storage (G') and loss (G") moduli of G3BP1 condensates measured using pMOT.** The complex moduli were measured after (a) 4 hr, (b) 6 hr, (c) 9 hr, (d) 12 hr, (e) 16 hr, and (f) 18 hr of sample age. For each sample, three independent replicates are shown here. (g) Complex moduli corresponding to all the condensate ages shown in (a)-(f) scaled along the frequency axis using scaling factor as the ratio of frequency $\omega$ and the respective crossover frequency $\omega_c$ at a sample age. The scaling shows a collapse of complex moduli corresponding to all the ages onto a single master curve depicting G3BP1 condensates obey time-ageing-time superposition.

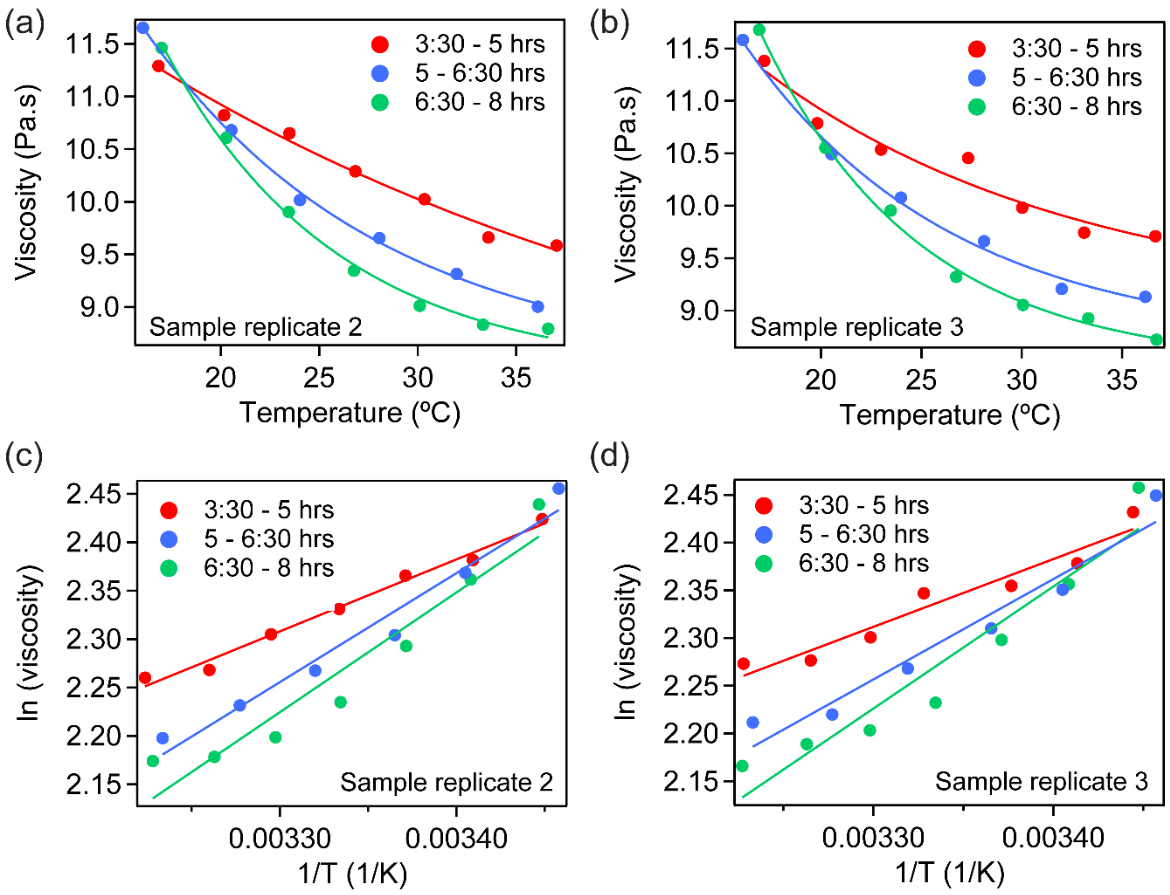


**Supplementary Figure 8. Viscosity of G3BP1 condensates as a function of temperature follows the Arrhenius law of flow activation energy.** Temperature-dependent viscosity of G3BP1 condensates, measured by temperature-controlled VPT nanorheology. Data were obtained for three independent sample preparations (Sample replicate 1–3). Sample replicate 1 is shown in Figure 2d-e in the main text. Here, the data are shown for sample replicate 2 (a, c) and sample replicate 3 (b, d). In (a) and (b), points are the data; solid lines are fits to the Arrhenius relation of viscous flow. (c) and (d) show linear fits of $ln(viscosity)$ vs $1/T$ for sample replicate 2 and sample replicate 3, respectively. The slopes of the lines correspond to the activation energy of network reconfiguration at different sample ages. The aggregated data from all three samples were used to estimate the activation energy values shown in Figure 2f.

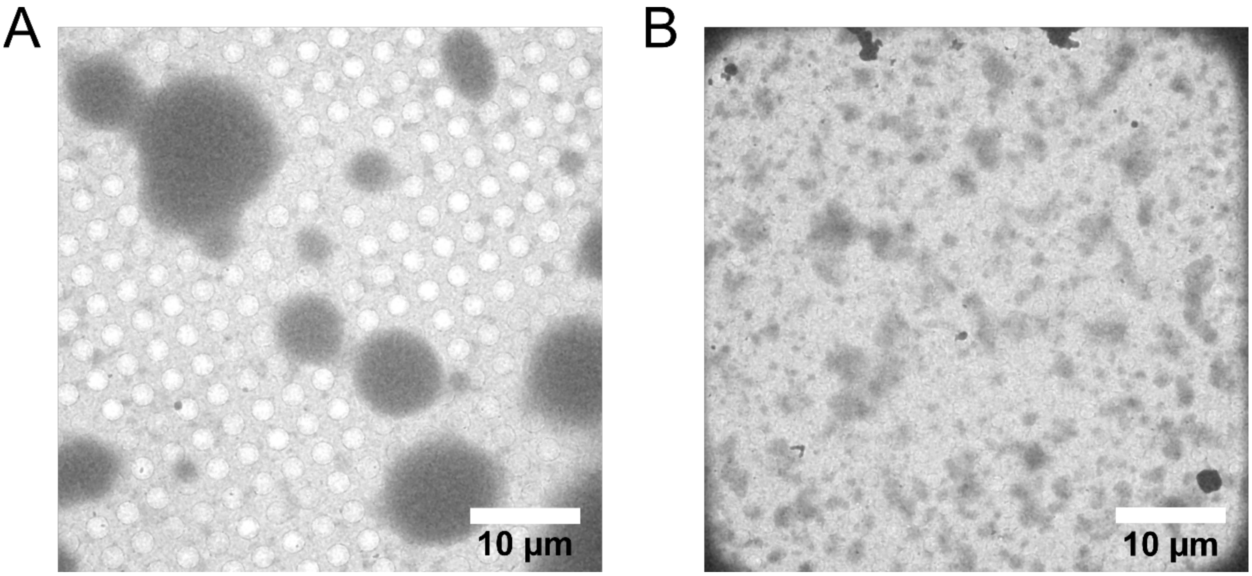


**Supplementary Figure 9. Lower-magnification cryo-electron micrographs at different sample ages** (A) Overview image of G3BP1 condensates prepared freshly (0 hr). Condensates appear round and spherical as expected. (B) Overview image of 36 hr aged G3BP1 condensates. Condensates display irregular shapes likely due to condensate breakage during sample blotting and vitrification.

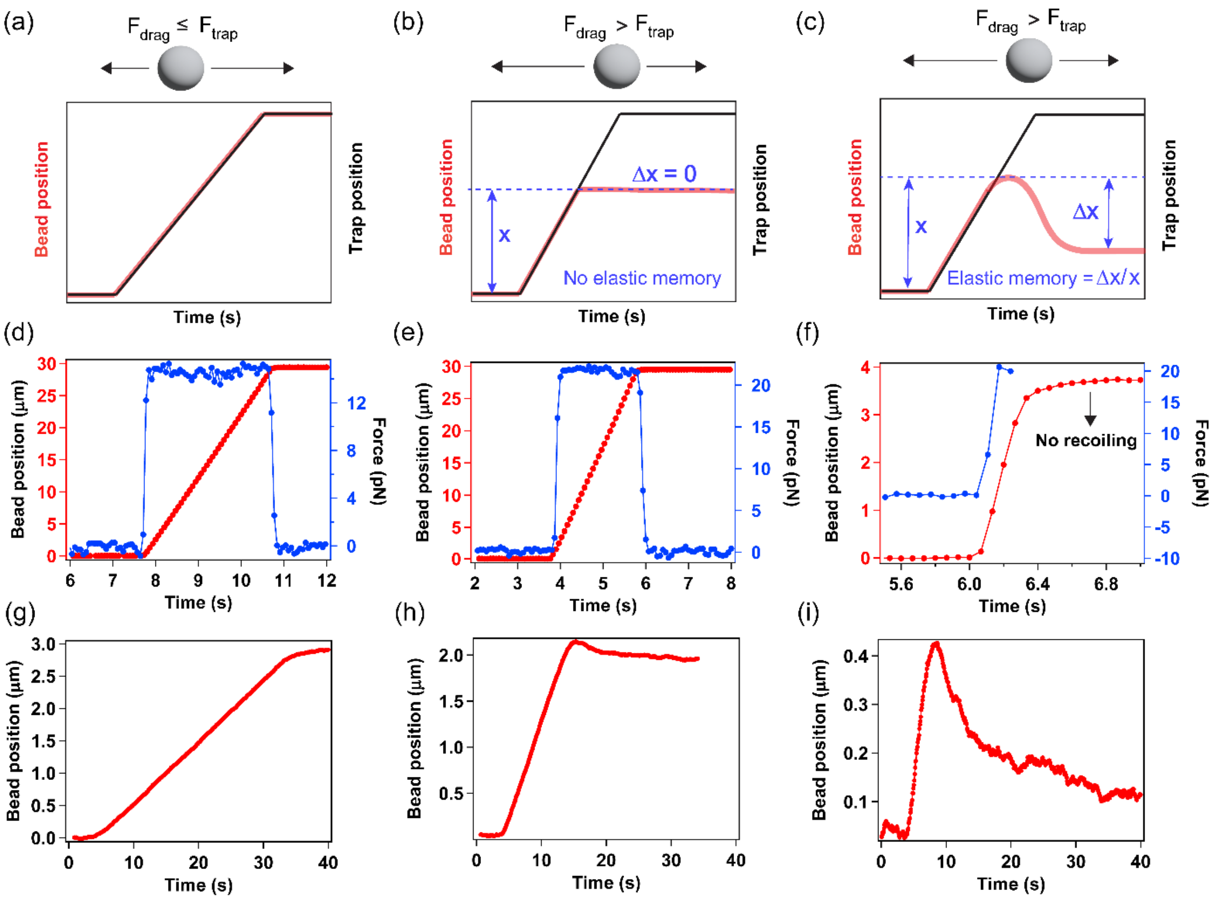


**Supplementary Figure 10. Laser tweezer-based creep test performed with 80% glycerol as a test sample and the effect of increased drag force on the bead position.** A bead of 2 μm in diameter was trapped in 80% glycerol solution. The optical trap was programmed to move to 30 μm with three different constant velocities (a) 10 μm/s, (b) 15 μm/s, and (c) 20 μm/s. The bead trajectory (red) and the force experienced by the bead (blue) are plotted here at a fixed trap stiffness. The viscosity of the glycerol solution extracted from the Stokes Drag forces in (d) and (e) was 0.072±0.005 Pa.s and 0.073±0.002 Pa.s, respectively. These viscosity values are close to the previously reported value of 0.066 Pa.s[2]. The viscosity for panel (f) was not determined because the bead was lost from the trap due to the viscous drag force on the bead exceeding the harmonic restoring force of the trap at this velocity. Notably, the bead does not recoil after it is released from the trap, indicating purely dissipative behavior of the fluid. For panels (g)-(i), a bead of 1 μm in diameter was trapped inside G3BP1 condensates after 12:30 hr of sample age. The optical trap was programmed to move to 3 μm at three constant velocities: (g) 0.1 μm/s, (h) 0.5 μm/s, and (i) 1 μm/s.

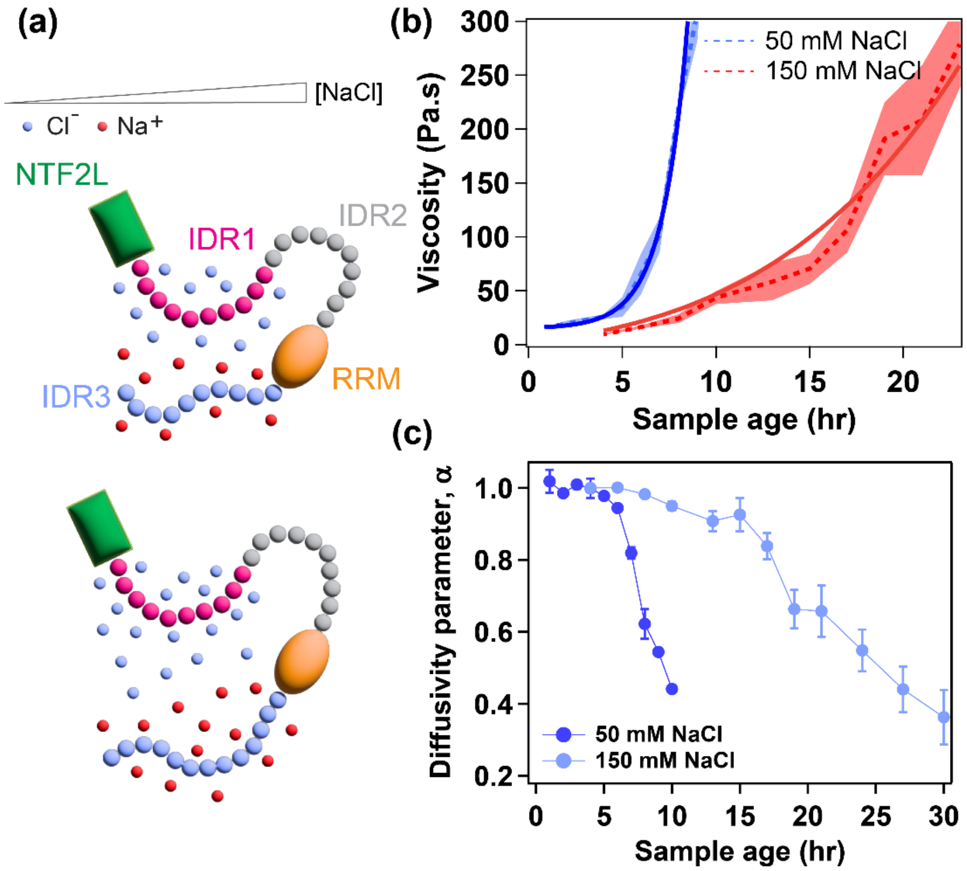


**Supplementary Figure 11. Role of electrostatic screening on homotypic G3BP1 condensate ageing.** (a) Schematic representation of the effects of monovalent salt on the homotypic interactions between IDR1 and IDR3 of G3BP1 molecules. (b)-(c) Variation of viscosity and diffusivity parameter as a function of sample age at different salt concentrations: 50 mM and 150 mM NaCl. Lower salt concentration mediates faster ageing of G3BP1 condensate.

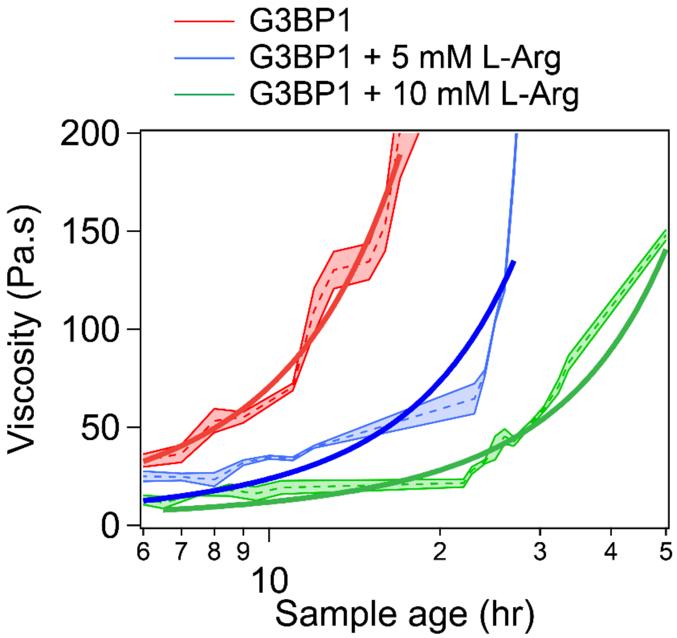


**Supplementary Figure 12. Role for arginine-mediated buffering of homotypic interactions that drive G3BP1 condensate ageing.** Variation of viscosity as a function of sample age at 5 mM and 10 mM L-Arg concentrations. These results show that L-Arg delays G3BP1 condensate dynamical arrest in a dose-dependent manner.

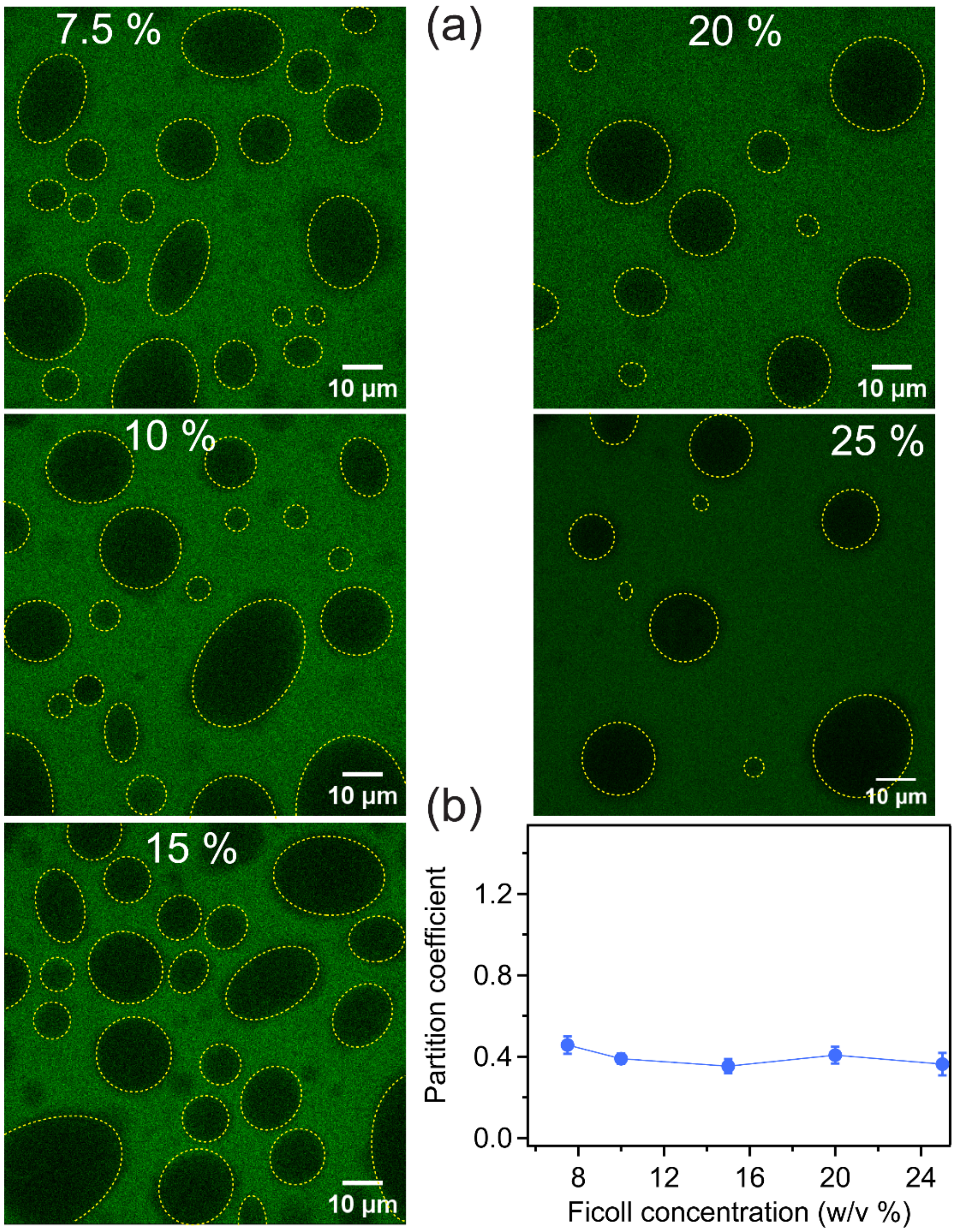


**Supplementary Figure 13. Ficoll-70 promotes the formation of heterotypic G3BP1-RNA condensates**. (a) Partitioning of Fluorescein-labeled Ficoll-70 inside G3BP1-RNA condensates at different concentrations of Ficoll-70, images shown for 7.5%, 10%, 15%, 20%, and 25% (w/v) Ficoll-70 concentrations. (b) Quantification of the partition coefficient of Ficoll-70 inside the G3BP1-RNA condensates.

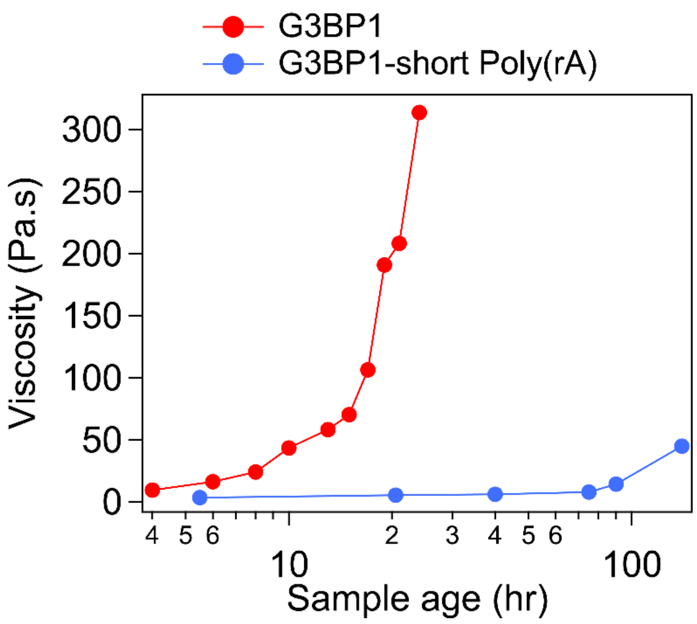


**Supplementary Figure 14. Effect of Poly(A) RNA on dynamical arrest of G3BP1 condensates at 150 mM NaCl condition**. No significant changes in condensate viscosity were observed even after 5 days of condensate age.

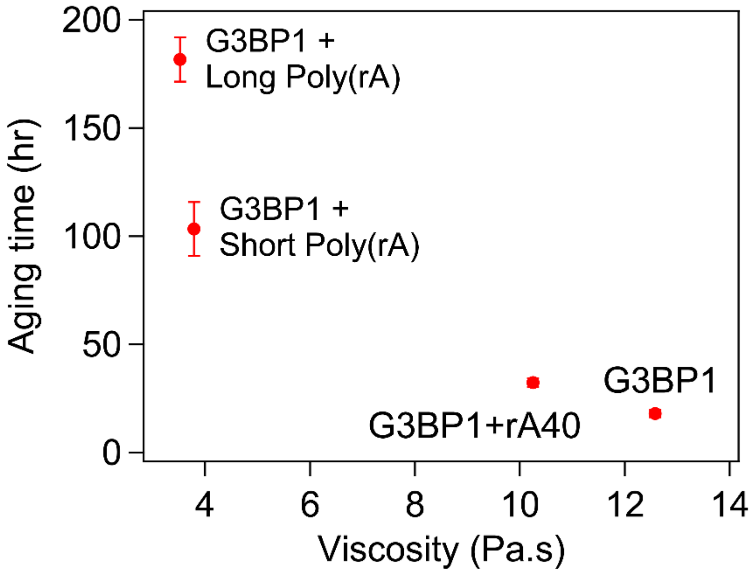


**Supplementary Figure 15. Correlation between ageing timescale and viscosity of nascent G3BP1-RNA condensates.** Ageing timescale inversely correlates with the viscosity of the nascent condensates, with lower viscosity corresponding to a longer ageing timescale of G3BP1-RNA condensates.

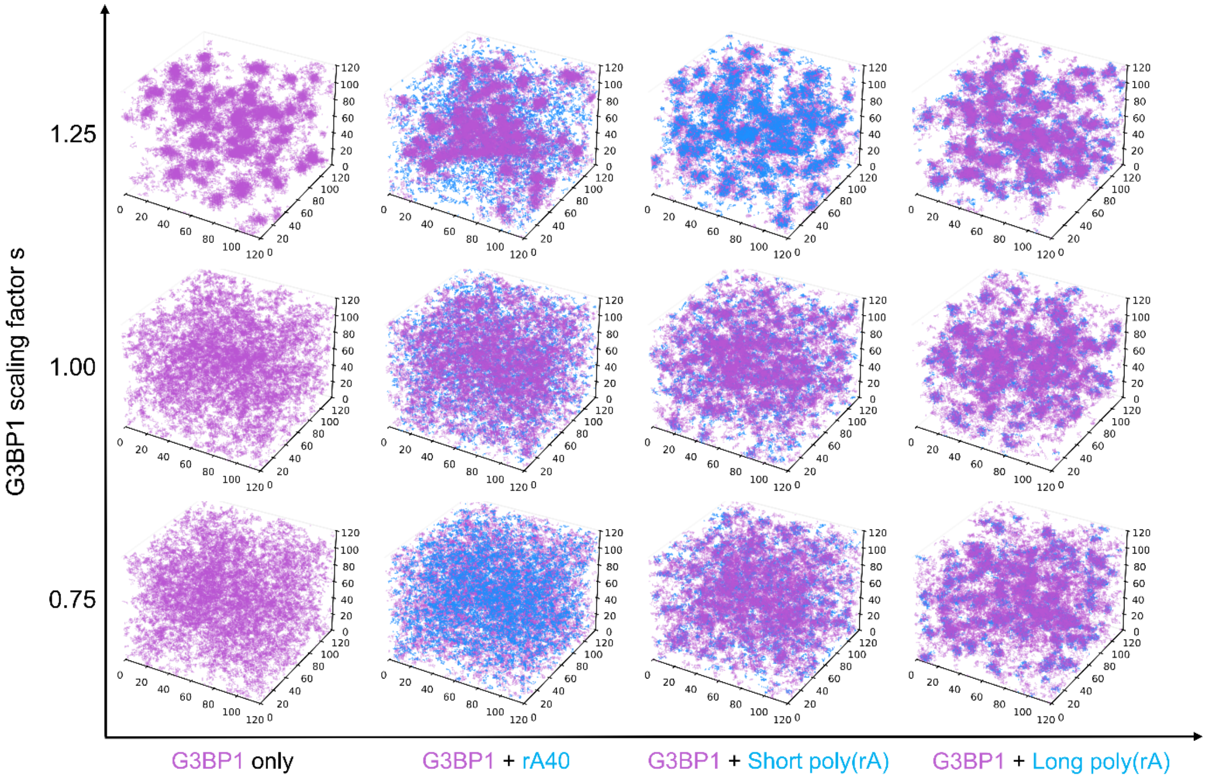


**Supplementary Figure 16. Absence of macrophase separation of G3BP1 clusters in the presence of RNA.** (a) Simulation snapshots showing G3BP1-RNA clusters at different scaling factors, $s$=0.75, 1.00, and 1.25 for rA40, short Poly(rA), and long Poly(rA) RNA. For all the RNA and interaction scaling factors, RNA promotes the association of G3BP1-RNA clusters without the formation of macrophases.

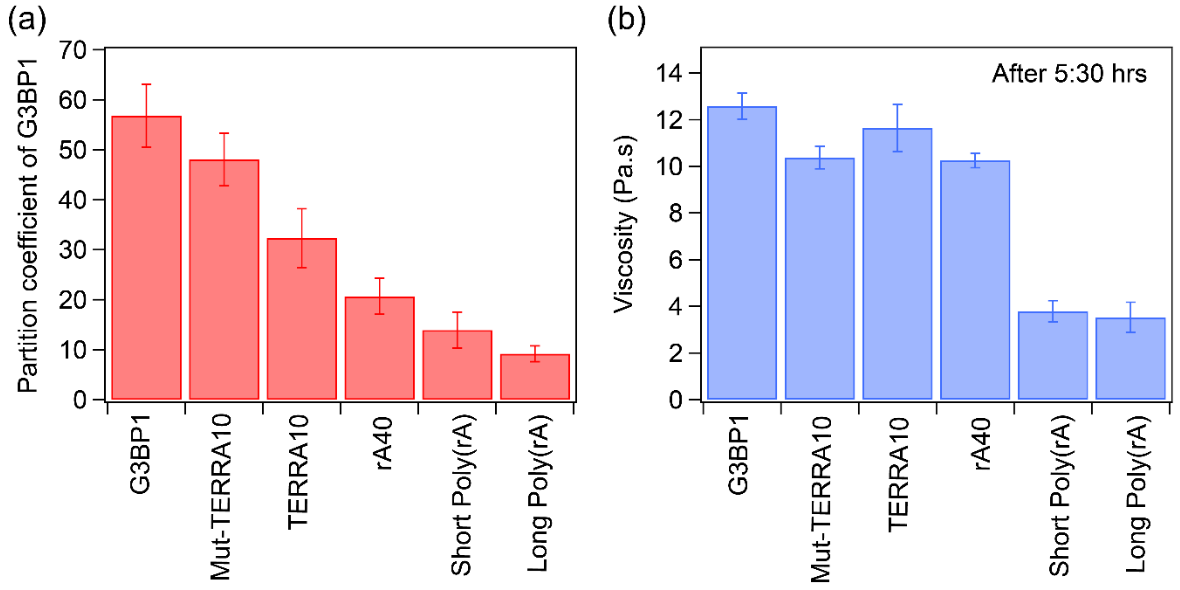


**Supplementary Figure 17. RNA regulates G3BP1 partitioning in the dense phase in a length and structure-specific manner.** (a) Partitioning of fluorescently labeled G3BP1 inside G3BP1-Ficoll and G3BP1-RNA condensates with different RNA substrates. The partitioning was quantified in terms of partition coefficients from confocal imaging of condensates with Atto488-labeled G3BP1. 200 nM Atto488-labeled G3BP1 was used for imaging. (b) Comparison of the viscosity of G3BP1-Ficoll and different G3BP1-RNA condensates after 5.5 hrs of sample preparation.

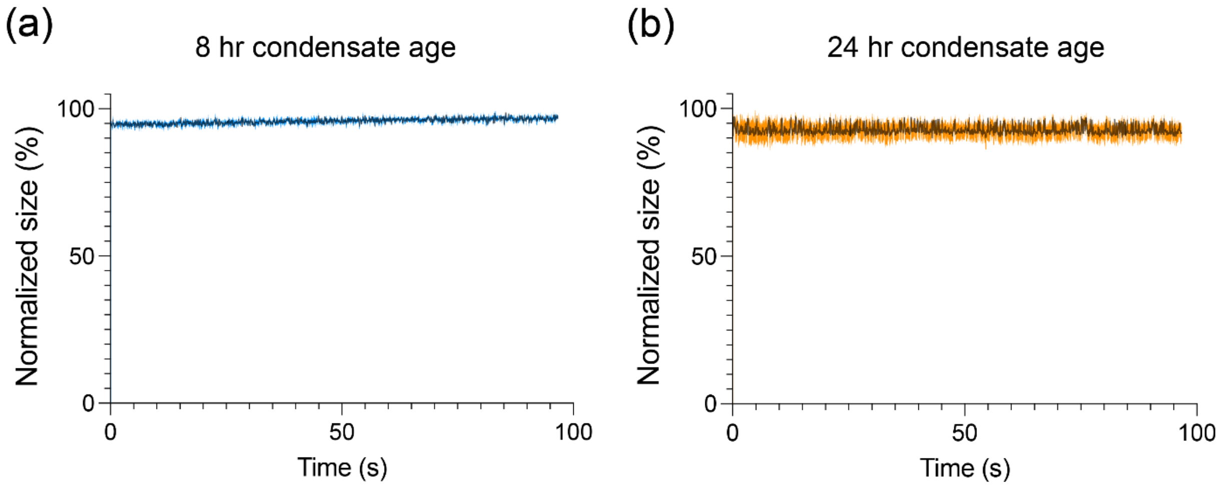


**Supplementary Figure 18. Stability of the G3BP1 condensates against flow-induced dissolution.** A single G3BP1 condensate at (a) 8 hr and (b) 24 hr sample age was optically trapped and moved to the buffer channel in a multi-channel microfluidic device. Under constant flow of the buffer, both (a) 8 hr and (b) 24 hr old G3BP1 condensates were resistant to dissolution, as measured by no significant change in their size. These persistent G3BP1 condensates were utilized to measure the efflux rate of poly(rA) RNA out of the condensate phase under flow at different sample ages.

**Supplementary Table 1**: Fitting parameters estimated from the fitting of the viscosity vs sample age data using the equation $\eta = \eta_0 exp\left[\left(\frac{t}{t_c}\right)^{\beta}\right]$ for G3BP1, G3BP1−RNA, G3BP1−Caprin-1, G3BP1−small molecule condensate systems.

| Condensate system | β | $t_c$ |
|---|---|---|
| G3BP1 (150 mM NaCl) | 0.44 ± 0.02 | 0.47 ± 0.11 |
| G3BP1-rA40 | 0.44 ± 0.05 | 1.03 ± 0.38 |
| G3BP1-short Poly(rA) | 0.78 ± 0.09 | 20.67 ± 4.09 |
| G3BP1-long Poly(rA) | 0.88 ± 0.15 | 53.89 ± 7.88 |
| G3BP1-TERRA10 | 0.44 ± 0.09 | 1.67 ± 0.94 |
| G3BP1-mut-TERRA10 | 0.53 ± 0.05 | 1.62 ± 0.41 |
| G3BP1-Caprin1 | 0.77 ± 0.08 | 3.42 ± 0.71 |
| G3BP1 (50 mM NaCl) | 0.37 ± 0.05 | 0.21 ± 0.12 |
| G3BP1- 5 mM L-arg | 0.43 ± 0.11 | 0.72 ± 0.68 |
| G3BP1- 10 mM L-arg | 0.42 ± 0.02 | 1.20 ± 0.22 |

### Supplementary Video Captions

**Video S1:** Representative video particle tracking movies of 200 nm yellow-green, fluorescent beads inside G3BP1 condensates after 4 hr, 10 hr, 24 hr, 27 hr, 29:30 hr, and 32:30 hr of sample age showing dynamical arrest of G3BP1 condensates with ageing.

**Video S2:** Fluorescence recovery after photobleaching (FRAP) of Atto488-labeled G3BP1 inside G3BP1 condensates after 5 hr and 23 hr of sample age.

**Video S3:** G3BP1 droplet fusion events after 30 min, 90 min, and 210 min of sample age showing age-dependent fusion dynamics of G3BP1 condensates.

**Video S4:** Real-time position of a 1 μm trapped bead inside 4 hr, 16 hr, and 23.5 hr old G3BP1 condensate with a constant velocity of 0.5 μm/s. These active microrheology (aMOT) measurements reveal age-dependent emergence of elastic memory in G3BP1 condensate network.

**Video S5:** Representative videos for early age (8 hr after sample preparation) and aged (24 hr after sample preparation) G3BP1-RNA condensates showing the efflux of labeled RNA molecules from G3BP1-RNA condensates as change in RNA intensity with time.

### Methods and materials

### Materials

**Purification of G3BP1 protein.** Results on G3BP1 condensate ageing shown in Figures 1-2 and Supplementary Figures 1-4 were obtained utilizing purified G3BP1 prepared in Dr. J. Paul Taylor's lab at St. Jude Children's Research Hospital. Briefly, GST-tagged G3BP1 protein was expressed and purified from E. coli BL21 (DE3) cells as reported before[3]. The GST tag was removed via TEV cleavage. SDS-PAGE was performed to analyze the purity of recombinant G3BP1 protein and was stored at -80 °C for further use.

The results on condensate ageing shown in Figure 3-6 and supplementary Figure 5-20 were obtained on G3BP1 purified in the Banerjee lab. The construct of full-length G3BP1 was synthesized and cloned into a pET His6 MBP N10 TEV LIC cloning vector (Addgene plasmid #29706 by GenScript USA Inc. (NJ, USA)). The MBP-His-G3BP1 construct was transformed into BL21-DE3 pLysS cells, which were grown in LB media at 37 °C to an OD600 of ~0.7, then induced with 0.6 mM IPTG and incubated overnight at 16 °C. Cells were harvested by centrifugation at 4000 g for 30 mins and lysed with an ultrasonicator in lysis buffer containing 50 mM Tris-HCl (pH 7.5), 500 mM NaCl, protease inhibitor cocktail (Thermo Fisher Scientific), 2 mM DTT, 1 mM phenylmethylsulfonyl fluoride (PMSF, Sigma-Aldrich), and 10 mM imidazole. After centrifugation at 37,500 x g for 30 min at 4 °C, the extracted supernatant was then incubated with Ni-NTA resin (pre-equilibrated with the lysis buffer) overnight at 4 °C. The resin was then washed with 1 M NaCl containing wash buffer and subsequently eluted with 50 mM Tris-Cl pH 7.5, 300 mM NaCl, 2 mM DTT, and 500 mM imidazole-containing buffer.

The combined elution fractions were then passed through a HiRes Capto Q 5/50-Anion exchange column (Cytiva) and washed with a buffer containing 50 mM Tris-Cl pH 7.5, 2 mM DTT, 50 mM NaCl. A step gradient elution from 50 mM to 1 M NaCl in 50 mM Tris-Cl pH 7.5 and 2 mM DTT-containing buffer. Elution fractions containing G3BP1 were combined and incubated overnight at 4 °C with TEV protease (purified in-house). The cleaved His-MBP tag was removed by subsequent incubation with Ni-NTA resin, and the untagged G3BP1 in the supernatant was collected. The purified protein was then buffer exchanged into 50 mM Tris-Cl pH 7.5, 400 mM NaCl, and 1 mM DTT, using a Zeba spin column (Thermo Fisher Scientific). The final sample was then concentrated and stored at -80 °C for future use. Fluorescence labeling of G3BP1 with Atto488 was achieved through Cys-maleamide chemistry following the manufacturer-provided instructions (Invitrogen).

Both G3BP1 preparations were rigorously characterized, and their condensates exhibited consistent viscosities and aging timescales.

**Caprin-1 protein**. Caprin-1 protein was received as a gift from Dr. J. Paul Taylor's lab at St. Jude Children's Research Hospital. The purity of the protein was assessed using liquid chromatography mass spectrometry (LC-MS) (see Supplementary Figure 19; theoretical mass of Caprin-1 is 78,372 Da).

The protein was acidified and diluted using 5% formic acid before LC-MS measurements. LC-MS was performed on a Thermo Scientific Orbitrap Exploris 480 mass spectrometer coupled to an UltiMate 3000 nanoLC system (Thermo Fisher Scientific). Samples were desalted online and separated on a nanoEase M/Z Protein BEH C4 column (Waters; 300 Å, 5 µm, 300 µm × 50 mm) before mass spectrometric analysis. Mobile phase A consisted of 0.2% formic acid in water, and mobile phase B consisted of 0.2% formic acid in acetonitrile. Peptides were eluted using a 5-min gradient. Mass spectrometric data were acquired in positive ion mode using a full MS scan over an m/z range of 400–4000 at a resolution of 60,000. The resulting raw data were processed using Thermo Scientific BioPharma Finder (version 5.0) software. Deconvolution was performed using the Xtract algorithm to determine the isotopically resolved mass of the protein.

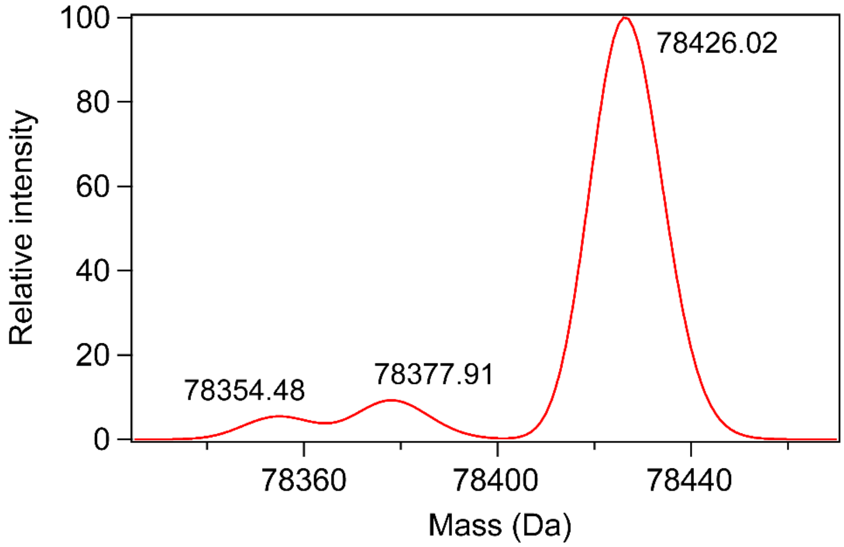


**Supplementary Figure 19.** Mass spectrum of Caprin-1 protein determined using LC-MS, showing the measured mass of Caprin-1 protein as 78,426 Da.

**RNA stocks.** rA40, TERRA10, and mut-TERRA10 RNA were purchased from Integrated DNA Technologies (IDT). Short poly(A) (300–4400 nucleotides; Catalog #P9403) and long poly(A) (2,100–10,000 nucleotides; Catalog #10108626001) RNAs were purchased from Sigma-Aldrich. The RNAs were reconstituted in RNase-free water followed by centrifugation at 23,000 × g for 2 min to remove any solid particles. The supernatant was extracted with a final concentration of at least 5 mg/ml, which was subsequently stored at −20 °C. The resulting stock solutions were divided into multiple aliquots before storage. We further confirmed the absence of microscale aggregates in all RNA stock solutions through light microscopy.

**Microrheology probes**. 200 nm yellow-green carboxylate-modified polystyrene beads (FluoSpheres, Invitrogen; Catalog #F8811) were used for VPT measurements. For pMOT and active microrheology measurements, 1 μm yellow-green carboxylate-modified polystyrene beads (FluoSpheres, Invitrogen; Catalog #F8823) were used for G3BP1 condensates and 2 μm polystyrene beads (FluoSpheres, Invitrogen; Catalog #F8824) were used for control experiments with glycerol. The bead stocks were diluted to have 0.0002% solids in the final sample volume for VPT nanorheology and 0.002% for pMOT and active microrheology measurements.

**Methods**

**G3BP1 phase separation and sample chamber preparation**

G3BP1 at 50 μM was added to the buffer containing 50 mM HEPES pH 7.5, 150 mM NaCl, and 0.2 mM DTT. Ficoll-70 at 10 % (w/v) was mixed to induce phase separation. For imaging, 200 nM Atto488-labeled G3BP1 was mixed with the unlabeled G3BP1. For rheology experiments, probe particles were added to the mixture and were well mixed before inducing phase separation. G3BP1 condensate samples were placed on an 18 x 18 mm Tween20 20% (v/v) coated microscope coverslip and sandwiched with a 75 x 25 x 1 mm thick microscope glass slide using three layers of double-sided tape (Scotch 3M). Mineral oil was then injected into the chamber surrounding the sample from all directions to prevent sample evaporation over time.

**Fusion of suspended droplets using optical traps**

Droplet fusion events were conducted using optical traps to investigate the fusion timescale of G3BP1 condensates, as previously described[4]. Briefly, sample chamber containing G3BP1 condensates were prepared as described in "G3BP1 phase separation and sample chamber preparation section". Condensates were trapped using a dual-trap optical tweezer system (LUMICKS, C-trap) far from each other in two optical traps (using a 1064 nm laser) and then brought into proximity. Trap 2 was held at a fixed position while Trap 1 was programmed to move at a constant speed of 40-100 nm/s in the direction of Trap 2. Coalescence started upon contact due to interfacial tension. The motion of the Trap 1 was stopped when the fused droplet relaxed to a spherical shape for the nascent condensates or stopped relaxing in the case of aged condensates.

**Fluorescence Recovery after Photobleaching (FRAP)**

Samples were prepared as described in "G3BP1 phase separation and sample chamber preparation" section. The sample chamber was then placed on a confocal microscope (LUMICKS, C-trap). For all FRAP experiments, the bleaching region to 0.6 x 0.6 µm was fixed to ensure that the changes in the FRAP recovery time are only due to the probe dynamics and not due changes in the bleaching area[5-7]. For each sample, we measured the FRAP traces for the nascent and aged condensates. The recovery traces were corrected for photofading and normalized with respect to the bleaching depth following previously published procedures[4]. The recovery traces were then averaged, and the error was estimated as the standard deviation at each time point. To estimate the recovery half-time, we fitted individual recovery traces using the following equation[4,5]

$$I_{recovery}(t) = \frac{I_0 + I_\infty \frac{t}{\tau_{1/2}}}{1 + \frac{t}{\tau_{\frac{1}{2}}}} \tag{1}$$

Here, $I_0$, $I_\infty$, and $\tau_{1/2}$ are fitting parameters; $\tau_{1/2}$ represents the FRAP recovery half-time. The mobile fraction was estimated from the fitting. The reported mobile fraction was taken as the average of the values extracted from the fits of individual FRAP measurements. The error was estimated as the standard deviation from the mean values.

**Video particle tracking (VPT) nanorheology**

Video particle tracking (VPT) measurements were performed on phase-separated G3BP1 condensates prepared as described in the "G3BP1 phase separation and sample chamber preparation" section. Yellow–green carboxylate-modified polystyrene beads of 0.2 µm size were mixed during sample preparation. These beads were confirmed to partition into the dense phases of phase-separated G3BP1 condensates and were used as probe particles for VPT measurements. The condensates were allowed to fuse before imaging and were rested on the microscope for 3 hr before data collection. The bead motion inside the G3BP1 condensates was captured at 24 °C in the form of videos using a 100× oil immersion objective lens and a Teledyne FLIR Blackfly S USB3 CMOS camera for 1000 frames with 100 ms exposure time. The acquired videos were processed using Fiji (version 1.54f). The probe particles were tracked to get their trajectories using the Trackmate software plugin in Fiji[8]. During tracking, intensity and quality filters were used to avoid tracking aggregated probe particles.

**Temperature-controlled VPT**

G3BP1 condensate samples were sandwiched between a coverslip and a glass slide, similar to what is described in the "G3BP1 phase separation and sample chamber preparation" section above. The sandwich was then placed on a custom-built thermal stage (INSTEC) which is attached to a Zeiss Primovert inverted microscope with a 100x oil-immersion objective lens. Teledyne FLIR Blackfly S USB3 CMOS camera was used for imaging. Condensates were allowed to settle on the glass slide for ~3 hr prior to the start of the measurement. Each condensate contained about 50-80 fluorescent microspheres (200 nm). The temperature stage was first set to the lower experimental temperature (10 °C). Due to the objective acting as a heat sink, the actual temperature within the sample was different from the preset temperature of the thermal stage. To obtain an accurate measurement of the sample temperature, we used a thermocouple with a heat insert that touches the glass directly inside the thermal stage chamber on the edge of the condensate sample. Within 10-20 minutes, the temperature reading of the thermocouple was observed to be equilibrating around ~15 °C (preset temperature = 10 °C). To start imaging, the microscope objective was focused on a particular condensate. A time-lapse video was collected for 1000 frames at a rate of 10 frames per second (exposure time was set to 100 ms). The temperature was increased in steps of 3 °C, and the sample was left to equilibrate for 15 minutes at each temperature. Then, a similar time-lapse video of the same condensate was collected to record the motion of the particles at the new temperature. This procedure was repeated to collect 7 points between 15 °C and 37 °C (preset temperature = 40 °C). Three trials were done for three independent sample preparations.

**Data analysis: Temperature-controlled VPT**

Movies of condensates containing diffusing probe particles at different temperatures were processed first using Fiji-ImageJ software[9]. Each movie contained 1000 frames. The particles were tracked using the TrackMate software plug-in in Fiji[8]. In particle tracking, intensity filters were used to exclude any particle aggregates from tracking. The extracted trajectories were corrected for drifting by subtracting the center of mass trajectory, which was calculated using the velocities of particles in the following way

$$\boldsymbol{X_{COM}}(t) = \boldsymbol{X_{COM}}(k\Delta t) = \boldsymbol{X_0} + \sum_{k=0}^{k\Delta t} \frac{1}{N} \sum_{i=1}^{N} \boldsymbol{v_i} \Delta t \tag{2}$$

Where $k$ is the frame number, $\Delta t$ is the frame time, and $N$ is the number of particles per frame. $\boldsymbol{X_0}$ is the center of mass vector of the first frame. After correcting the trajectories for drifting, the mean squared displacement (MSD) was calculated as

$$MSD(m\Delta t) = \frac{1}{\Delta t} \frac{1}{N-m} \sum_{i=1}^{N-m} (x_{i+m} - x_i)^2 + (y_{i+m} - y_i)^2 \tag{3}$$

Where $m$ is the lag time in frames. The MSD was then fitted using[10,11]

$$MSD(\tau) = 4D\tau^{\alpha} + N \tag{4}$$

To obtain the diffusion coefficient $D$ of the particles. For all the systems, we insured that the value of the diffusivity exponent $\alpha$ is equal to 1 by choosing an appropriately long frame time to ensure measuring the terminal viscous behavior. The diffusion coefficient is then converted to viscosity using the Stokes-Einstein equation[10,11]

$$\eta = \frac{k_B T}{6\pi DR} \tag{5}$$

Where $R$ is the radius of the particles and $T$ is the temperature of the sample.

To estimate the activation energy of the G3BP1 condensates at different sample ages the movies collected at different temperatures were analyzed as described above and the value of the viscosity was plotted against the temperature. The viscosity variation as a function of temperature followed Arrhenius law of viscous flow,

$$\eta = \eta_0 exp\left(\frac{E_A}{RT}\right) \tag{6}$$

Further, In $\eta$ against $1/T$ was plotted and was fitted with

$$\ln\eta = \ln\eta_0 + \frac{E_A}{R}\left(\frac{1}{T}\right) \tag{7}$$

to estimate the activation energy of the G3BP1 condensates at different ages. The activation energy was estimated for three sample ages for three independently prepared samples. The average value of the activation energy is reported, and the error is estimated as standard deviation from the mean.

**Passive microrheology with optical tweezers (pMOT)**

G3BP1 condensate samples were sandwiched between a coverslip and a glass slide, similar to what is described in the "G3BP1 phase separation and sample chamber preparation" section above. pMOT experiments were performed following our previously published protocol[10]. The sample was loaded onto a correlative optical-tweezer and confocal microscopy setup (LUMICKS, Ctrap). The sample was equilibrated for ~3 hrs until all droplets had settled on the coverslip surface and no fusion event was observed. The optical trap was used to trap a bead within a condensate at minimal power (~10-50 μW). The trapped bead was then tracked utilizing a brightfield camera at 500 Hz using a template-matching algorithm for 10-30 minutes at different sample ages. For determination of nanocages, the tracked bead trajectories at different sample ages were further analyzed to estimate the dimensions of nanocages from the probability distribution of the bead positions.

**Data analysis: pMOT**

The two-dimensional trajectories of the trapped particles were analyzed to calculate the complex modulus of the condensate as a function of frequency $\omega$

$$G^*(\omega) = G'(\omega) + i\,G''(\omega) \tag{8}$$

Where $G'$ and $G''$ are the frequency-dependent elastic and viscous moduli, respectively. Briefly, the trajectories are first detrended using a spline-based detrending algorithm to remove the long-time drift of the trajectories. Next, each component (X and Y) of the trajectory is treated separately. For each one-dimensional trajectory (X or Y), we obtain the trap stiffness using the equipartition theorem[12-14]

$$\kappa_x = k_B T / \langle x^2 \rangle \tag{9}$$

Where $\kappa_x$ is the optical trap stiffness in the $x$ direction, $T$ is the temperature in Kelvin, and $k_B$ is the Boltzmann constant. Next, a normalized position autocorrelation function $g(\tau)$ is calculated. The autocorrelation function value at $\tau = 0$ is extrapolated using a spline algorithm. The autocorrelation function is then transformed into the frequency domain $\hat{g}(\omega)$ using a discrete Fourier transform algorithm[12-14]

$$-\omega^2 \hat{g}(\omega) = i\omega g(0) + \frac{\left(1 - e^{-i\omega t_1}\right)\left(g_1 - g(0)\right)}{t_1} + \dot{g}(\infty) e^{-i\omega t_N} + \sum_{k=2}^{N} \left(\frac{g_k - g_{k-1}}{t_k - t_{k-1}}\right)\left(e^{-i\omega t_{k-1}} - e^{-i\omega t_k}\right) \tag{10}$$

Lastly, the complex modulus is calculated using[12-14]

$$G^*(\omega) = G'(\omega) + i\,G''(\omega) = \frac{\kappa}{6\pi a}\left(\frac{i\omega \hat{g}(\omega)}{1 - i\omega \hat{g}(\omega)}\right) \tag{11}$$

Where $a$ is the radius of the particle (0.5 μm). For each trial, we extract two sets of $G'$ and $G''$ values for each frequency. For initial characterization of the nascent G3BP1 condensates (sample age ~4 hr), measurements over different samples were performed, and at each frequency, we average the values of $G'$ and $G''$ and report the final value. The error is calculated as the standard deviation from the mean. Further details of the experiment and data analysis, including various controls for the effects of condensate surface and glass surface on the measurement, are reported in our earlier work[10,12].

### Cryo-EM sample preparation and cryo-electron tomography

G3BP1 condensates were prepared as described above and measurements were performed for two sample ages. Nascent G3BP1 condensates (sample age - 0 hr) samples were vitrified immediately after condensate formation, whereas aged condensates (sample age – 36 hr) were obtained by allowing condensates to age for 36 hr before vitrification.

For both conditions, condensates were applied to glow-discharged Quantifoil Cu 1.2/1.3, 300-mesh grids. Grids were vitrified using a Vitrobot Mark IV (Thermo Fisher Scientific) with a blotting force of 1 and a 1 s blot time at 4 °C and 100% humidity, followed by plunge freezing into liquid ethane. Grids were stored in liquid nitrogen until imaging.

### Cryo-ET data acquisition

Cryogenic electron tomography was performed on a Titan Krios G4 (Thermo Fisher Scientific) operating at 300 kV at the Electron Imaging Center for Nanosystems (EICN), UCLA. Tilt series were collected using eTomo in a dose-symmetric scheme from –48° to +48° with 3° increments, maintaining a total cumulative electron dose below ~160 $e^-/Å^2$.

**Tomogram reconstruction and analysis**

Tilt series were aligned and reconstructed using weighted back-projection workflows in eTomo, and CryoFlow reconstructions were used for visualization and downstream analysis. Tomograms were examined and segmented in IMOD, and quantitative measurements, including FFT analysis and azimuthal averaging, were carried out in IMOD and ImageJ unless otherwise noted.

**Active microrheology using optical tweezers (aMOT)**

For the controlled experiments in 80% glycerol solution, a bead of 2 μm in diameter was trapped, and the optical trap was programmed to move to 30 μm with three different constant velocities (a) 10 μm/s, (b) 15 μm/s, and (c) 20 μm/s. For the active microrheology measurements inside G3BP1 condensates at different sample ages, 1 μm bead was trapped and moved to 3 μm inside the G3BP1 condensate with a constant velocity of 0.5 μm/s unless otherwise noted. Real-time position of the trapped bead after 4 hr, 12:30 hr, 16 hr, 21 hr, and 23.5 hr of the sample preparation are reported.

**Atomistic simulations**

We performed atomistic simulations to extract pairwise inter-domain interaction coefficients, which serve as proxies for second virial coefficients, between the folded domains and intrinsically disordered regions of G3BP1.

All-atom Monte Carlo simulations were performed using the ABSINTH implicit solvent model and forcefield paradigm as made available in the CAMPARI simulation package (http://campari.sourceforge.net)[15,16]. All simulations were performed using version 4.0 of the CAMPARI simulation package and the abs_3.5opls.prm parameter set, set in conjunction with optimized parameters for neutralizing excess $Na^+$ and $Cl^-$ ions[17]. Each of the simulations were performed using spherical droplets with a radius of 200 Å, which is over an order of magnitude larger than the dimensions of an individual domain, represents an ultra-dilute system for a pair of domains and includes explicit ions to mimic a concentration of 1.7 mM NaCl. Ensembles corresponding to a temperature of 340 K are used in the analysis reported in this work, which based on recent calibrations, has been shown to generate conformational statistics that are congruent with various experiments[18,19].

For simulations involving the folded domains, we used the predicted structure for G3BP1 from AlphaFold corresponding to the Uniprot ID Q13283[20]. An initial single simulation for each pair of domains was performed at a temperature of 20 K to build the IDRs and equilibrate the conformations of the RRMs to make sure the bond lengths and bond angles were consistent with the ABSINTH model. Then, the end PDB structure from these simulations was used as the input structure for ten independent simulations, each initiated using a distinct random seed. The backbone flexibility was limited by applying restraints on the starting structure, whereas the sidechain degrees of freedom were unrestrained. In a finite unbiased simulation, most of the inter-domain distances sampled would correspond to dissociated molecules. To overcome this problem, we applied an inter-domain distance restraint, modeled using a harmonic potential. Pairs of alpha carbon atoms located close to the centroid of each molecule were restrained to be within a shallow harmonic well, defined by a spring constant of 10 kcal/mol-Å and an equilibrium distance of 75 Å. Note that no restraints were applied on any other inter-residue or inter-atomic distances. Each simulation comprised a total of 4.0 x $10^7$ MC steps that combined the full spectrum of translational, rotational, pivot, local, and concerted moves. Conformations pertaining to the first 5.0 x $10^6$ MC steps were discarded as equilibration.

From the ensemble of equilibrated configurations for each of the distinct pairs of domains with ten independent simulations for each pair, we extracted statistics for the inter-residue distances using MDTraj[21] and SOURSOP[22]. These statistics were used to compute the ensemble average distances $\langle R_{XY}^{(ij)} \rangle$ between pairs of residues *i* and *j*, where the former is located on domain X and the latter on domain Y. To calibrate the observed pattern of inter-domain, inter-residue distances against a suitable prior, we performed reference simulations to model the expectations for an ideal, non-interacting pair of the domains of interest. In this non-interacting priority, the domains freely sample the available simulation volume, with the presence of the weak restraint as described above. These reference simulations were performed using the inverse power potentials introduced by Tran et al[23]. In these simulations, the non-bonded interactions are such that all terms, except the Lennard-Jones repulsions, were switched off. The exponent for the repulsive exponent was set to six, which corresponds to Maxwellian particles, and the ε was set to 0.0001. This choice was based on calibrations, which showed for IDRs, irrespective of the sequence, an exponent of six and ε of 0.0001, with all other non-bond interactions switched off, the ensemble-averaged intramolecular, internal distance profiles showed fractal behavior whereby the distances between pairs of residues *i* and *j*: $\langle R_{ij} \rangle \sim |j-i|^{0.5}$. This is the scaling expected for a Flory random coil[24-26]. Inter-domain, inter-residue distances extracted from simulations using pairs of domains based on the inverse-power potential with an exponent of six and ε of 0.0001 were used to extract ensemble-averaged distances denoted as $\langle R_{XY,\mathrm{ideal}}^{(ij)} \rangle$.

The effective inter-domain interaction coefficient between a pair of domains X and Y is described by a parameter $\Delta_{XY}$ (Also described in Shinn et al., 2025[27]). Values for $\Delta_{XY}$ are normalized and signed such that $-1 \leq \Delta_{XY} \leq +1$ by the number of residues in the respective domains. Values of $\Delta_{XY}$ less than 0 signify attractive interactions whereas $\Delta_{XY}$ values greater than 0 signify repulsive interactions. Values of $\Delta_{XY}$ that are near 0 signify ideal interactions. The magnitude of $\Delta_{XY}$ describes the extent of the interaction strength. The inter-domain interaction coefficients, $\Delta_{XY}$, are mapped to the sequence architecture of G3BP1 to generate the inter-domain interaction matrix.

**Coarse-grained simulations**

We employed a coarse-grained phenomenological G3BP1 model parameterized from all-atom simulations to uncover the intrinsic phase behavior and organization using LaSSI, a lattice-based Monte Carlo simulation engine.

Simulations were performed using LaSSI, a lattice-based Monte Carlo simulation engine that is based on the generalized bond fluctuation model[28]. Monte Carlo moves are accepted or rejected based on the Metropolis-Hastings criterion. Each simulation consisted of 2,500 G3BP1-mimicking chains and was performed on a cubic lattice with a box size of L = 120 lattice sites and periodic boundary conditions. Previous calibrations have shown that the number of molecules used in these simulations is adequate to avoid problems due to finite-size effects[29]. Each G3BP1 chain comprises five different bead types corresponding to each G3BP1 domain. The number of beads for each domain is determined by the average radius of gyration across an ensemble of equilibrated configurations obtained from the atomistic simulations involving the domain, and allocating one bead for every 6 Å. The pairwise interaction energies between each bead type are defined by the inter-domain interaction. In addition, a set of scaling factors is manually curated, and each scaling factor is uniformly applied to the interaction matrix. This procedure preserves the relative strengths of different interactions between different domain types while modulating the overall interaction strength. As a result, the balance between attractive and repulsive interactions is tuned without altering the relative interaction hierarchy. Each simulation comprises a total of 5.0 x $10^9$ Monte Carlo steps. Analyses were only performed using data from the second half of each simulation.

For the G3BP1-RNA simulations, we chose the appropriate chain lengths for each RNA-mimicking polymer based on the number of nucleotides. In each G3BP1-RNA system, the number of RNA molecules was determined such that the total number of RNA beads was equivalent and distributed in different lengths to investigate the effect of RNA length on G3BP1 phase behavior aligned to experimental conditions. The only attractive interactions between the RNA beads are with the sites of G3BP1 RRM1 and G3BP1 IDR3, whereas all other interactions with G3BP1 beads are set to be implicitly repulsive.

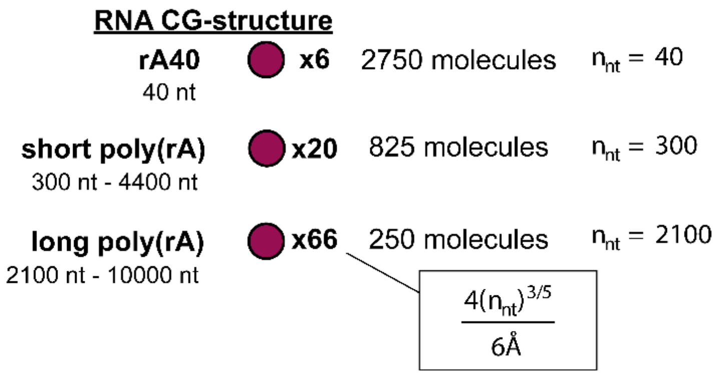


| | RNA | NTF | IDR1 | IDR2 | RRM | IDR3 |
|---|---|---|---|---|---|---|
| $\varepsilon_{RNA}$ | 0 | 0 | 0 | 0 | -0.25 | -0.25 |

**Supplementary Figure 20.** Coarse-grained structure of rA40, short Poly(rA), and long Poly(rA) designed for the G3BP1-RNA coarse-grained simulations. The number of nucleotides ($n_{nt}$) for the three RNAs was chosen based on the exact number for rA40 and lower end of the nucleotide range for short Poly(rA) and long Poly(rA) RNAs used in the experiments. The lower panel shows the interaction strength in the parameter $\varepsilon_{RNA}$ between RNA and G3BP1 interaction domains.

**Condensate efflux assay using correlative optical tweezers–fluorescence microscope system**

G3BP1 condensates containing trace amounts of poly(rA) with Cy5-dT$_{40}$ (for visualizing poly(rA)) at defined sample ages (either 8 hr or 24 hr) were flowed into a pre-equilibrated microfluidic device (μ-Flux setup, LUMICKS) integrated as part of the LUMICKS C-trap correlative optical tweezers–confocal fluorescence microscope system. Refer to the "G3BP1 phase separation and sample chamber preparation" section above for details on the condensate samples. The flow rate was set to ~1.2 bar to allow the flow of condensates into the designated flow chamber, with the trap power set to 10% (corresponding to ~100 μW; 1064 nm laser) to capture a condensate for efflux measurement. All operations were controlled using the Bluelake software package (LUMICKS). Once a condensate is trapped by the 1064 nm trapping laser, the flow rate is immediately reduced to 0.2 bar, and the condensate is moved to a waypoint in the adjacent channel, which contains the sample buffer alone. Next, continuous fluorescence microscopy (0.7 frames per second) with a Cy5 excitation laser line was conducted to monitor the fluorescence signal of Cy5-dT$_{40}$, serving as a proxy for the poly(rA) concentration within the G3BP1 condensate at either sample age (8 hr or 24 hr).

Analysis of the resulting fluorescence time-lapse videos was done using Fiji[30], as described previously[31,32]. Briefly, a circular ROI was used to measure the fluorescence intensity of the RNA partitioned to the condensate as a function of time using the 'Plot Z-axis profile' function of Fiji, which would serve as an estimate of RNA efflux rate. The measured intensity values in each measurement were normalized according to the maximum value. The fluorescence intensity as a function of time was plotted using GraphPad Prism 10, and the data were fitted with a single-phase decay model:

$$Y = Plateau + (Y_0 - Plateau) \cdot exp(-K \cdot X) \tag{12}$$

Using this relation, the time constant ($\tau$) is estimated as the inverse of the rate constant K, thereby providing a measure of the characteristic decay rate of the fluorescence signal from the RNA within a condensate. Such analyses were performed for n = 8 condensates for 8 hr time point and n = 6 for 24 hr time point, representing three independent replicates. The data in the final plot are shown as the mean $\pm$ SEM. For supplementary video 5, 8 hr and 24 hr time point videos were upscaled by 10x via bilinear interpolation to enhance clarity and improve visualization. Display range of the videos (i.e. contrast) were adjusted independently in 8 hr and 24 hr conditions to aid visualization.

**Condensate dissolution assay using a correlative optical tweezers–microscope system**

The experimental approach employed here is largely similar to that described in the '*Condensate efflux assay using correlative optical tweezers–fluorescence microscope system*' method section. The exception is the use of a higher frame rate brightfield camera (15 frames per second) to measure the whole condensate size as a function of time. The analysis pipeline used here is similar to that described in a previous report, with a few exceptions[32]. Briefly, the time-lapse brightfield videos were imported into Fiji[30], whereby the 'Find Edges' function was used to detect the droplet boundaries. Then, the 'Gaussian Blur' function was used to smooth the detected boundaries into a continuous region. Subsequently, thresholding was performed to binarize the image stack and thus distinguish the condensate from the background clearly. In some cases, the droplet interior (area within the detected droplet boundary) is not captured during the thresholding process owing to the innate bias of edge detection (from the step involving the 'Find Edges' function). To correct this, the 'Fill Holes' function is used to effectively capture the entire droplet area. Lastly, the 'Analyze Particles' function is used to measure individual droplet sizes as a function of time. The data were plotted using GraphPad Prism 10, reported as mean $\pm$ SEM, representing n = 3 condensates from three independent replicates.
